# scHiCcompare: an R package for differential analysis of single-cell Hi-C data

**DOI:** 10.1101/2024.11.06.622369

**Authors:** My Nguyen, Brydon P. G. Wall, J. Chuck Harrell, Mikhail G. Dozmorov

## Abstract

Changes in the three-dimensional (3D) structure of the human genome are key indicators of cancer and developmental disorders. Techniques like chromatin conformation capture (Hi-C) have been developed to study these global 3D structures, typically requiring millions of cells and an extremely high sequencing depth (around 1 billion reads per sample) for bulk Hi-C. In contrast, single-cell Hi-C (scHi-C) captures 3D structures at the individual cell level but faces significant data sparsity, marked by a high proportion of zeros. While differential analysis methods exist for bulk Hi-C data, they are limited for scHi-C data. To address this, we developed a method for differential scHi-C analysis, building on existing techniques in the HiCcompare R package. Our approach imputes sparse scHi-C data by considering genomic distances and creates pseudo-bulk Hi-C matrices by summing condition-specific data. The data are normalized using LOESS regression, and differential chromatin interactions are detected via Gaussian Mixture Model (GMM) clustering. Our workflow outperforms existing methods in identifying differential chromatin interactions across various genomic distances, fold changes, resolutions, and sample sizes in both simulated and experimental contexts. This allows for effective detection of cell type-specific differences in chromatin structure, which has meaningful associations with biological and epigenetic features. Our method is implemented in the scHiCcompare R package, available at https://github.com/dozmorovlab/scHiCcompare.

**Graphical abstract:** 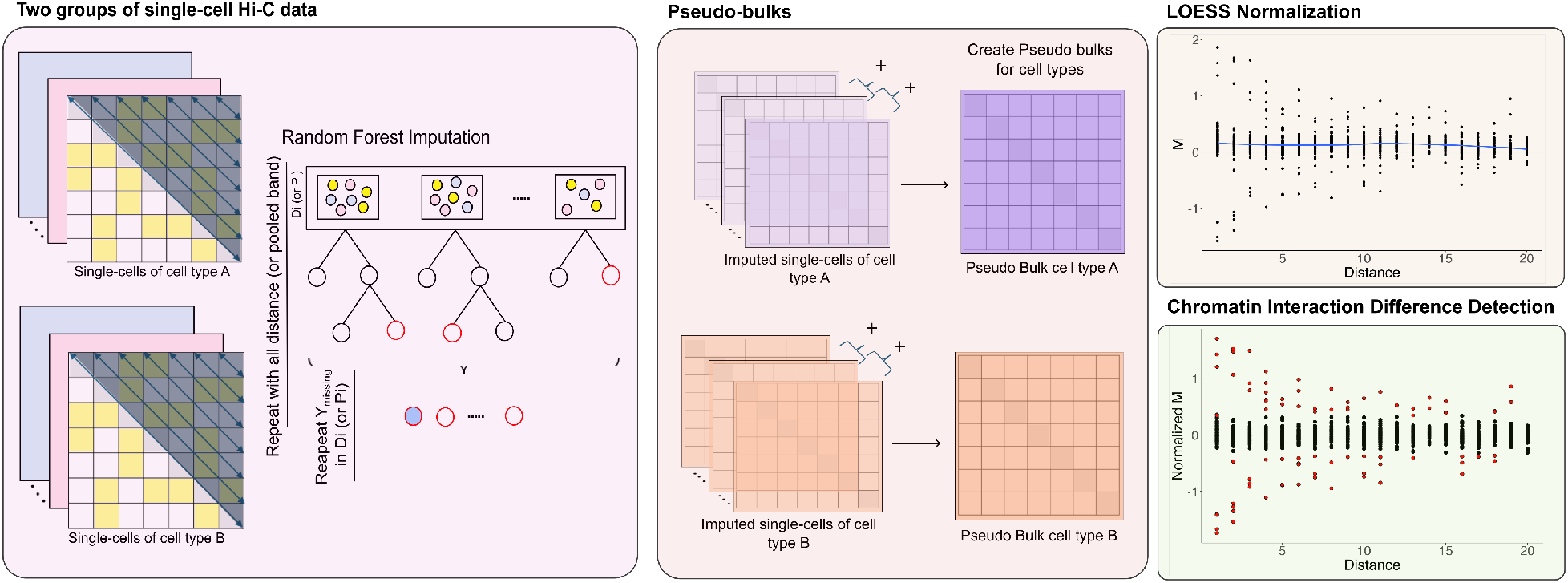

## Introduction

Many studies have demonstrated that the three-dimensional (3D) organization of chromosomes plays a fundamental role in regulating gene expression and maintaining genome stability [1–3]. Changes in the 3D structure of the genome are well-established hallmarks of cancer and developmental disorders, as such alterations can lead to gene dysregulation and contribute to disease progression [4]. Chromatin conformation capture technologies, such as Hi-C, have been developed to investigate the 3D structure of chromatin within the nucleus [5]. These technologies provide a comprehensive view of spatial interactions between different genomic regions, revealing how chromatin loops and domains contribute to genome organization and function. Chromatin conformation capture technologies have evolved not only to capture the 3D genome as an average across many cells, as seen with Hi-C, but also to examine chromosome and genome structures in individual cells through single-cell approaches [6].

Single-cell Hi-C (scHi-C) enables the mapping of 3D chromatin structures in single cells, offering insights into cell-cycle dynamics and cell type-specific chromatin configurations [6,7]. This emerging technology provides a more detailed understanding of chromatin organization and its variability across individual cells, revealing nuances that are not apparent in bulk analyses. However, it faces challenges such as high sparsity and significant heterogeneity in single-cell Hi-C data, similar to those encountered in single-cell RNA sequencing (scRNA-seq)—a technology that provides valuable insights into gene expression and cellular function at the single-cell level. Despite the availability of numerous differential analysis methods for RNA-seq and scRNA-seq, fewer methods are available for differential analysis in Hi-C data (e.g., HiCcompare [8], multiHiCcompare [9], Find [10]), and even fewer for scHi-C data (e.g., SnapHiC-D [11]).

We have developed a novel method to detect significant differential chromosome interactions (DCIs) between two chromosome-specific groups (pre-defined clusters) in single-cell Hi-C (scHi-C) data. To address the challenges of data sparsity, we apply random forest imputation to each scHi-C matrix, which accounts for variability among single cells and genomic distance heterogeneity. Although individual cells within a specific cell type exhibit some variability, we expect them to share similar chromatin interaction patterns characteristic of that cell type. Consequently, we convert these imputed single-cell data into group-specific pseudo-bulk Hi-C datasets, consolidating chromatin interaction information for each cell type. The improved HiCcompare method, which integrates normalization and differential detection, is then used to identify DCIs. Using simulated differences and experimental datasets, we demonstrate that scHiCcompare effectively and accurately identifies single-cell DCIs between two cell type groups. Our workflow also outperforms existing differential detection methods, such as SnapHiC-D [11] and scHiCDiff [12].

## Results

### Overview of differential analysis of scHi-C data using pseudo-bulk

Single-cell Hi-C (scHi-C) data allows for the identification of chromatin interactions at the individual cell level, enabling the detection of cell type-specific chromatin interaction patterns. While various methods exist for clustering scHi-C data, such as Higashi [13] and scHiCcluster [14], approaches for detecting differential chromatin interactions (DCI) between cell type-specific clusters remain limited. Existing methods like SnapHiC-D [11] and scHiCDiff [12] have attempted to provide solutions yet have notable shortcomings. SnapHiC-D [11] identifies cell type-specific chromatin contacts at a 10 kb resolution using a two-sided two-sample T-test, with missing data imputed by a random walk with restart algorithm and normalized using Z-scores. However, it lacks flexibility across different resolutions. Similarly, scHiCDiff [12] compares chromatin interaction bin pairs between two conditions using a Zero-Inflated Negative Binomial model, with imputation via a Gaussian convolution filter and normalization through scHiCNorm [15]. Despite its utility, scHiCDiff encounters challenges with chromosome-wide comparisons and scalability when analyzing high-resolution datasets.

To address these challenges, we developed scHiCcompare, a method specifically designed for differential chromatin interaction analysis across two pre-defined, chromosome-specific groups of scHi-C data. The scHiCcompare workflow has four main steps: (1) random forest imputation of missing data within individual genomic distance scHi-C data or pooled bands, accounting for the decay of interaction frequencies with distance, (2) pseudo-bulk matrix generation by summing interaction frequencies (IF) for individual cells within each cell type group, (3) joint normalization of the two pseudo-bulk matrices using locally estimated scatterplot smoothing (LOESS) [16] regression, and (4) differential chromatin interaction detection between matrices utilizing Generalized Mixture Model (GMM) clustering (Figure 1).

**Figure 1.**
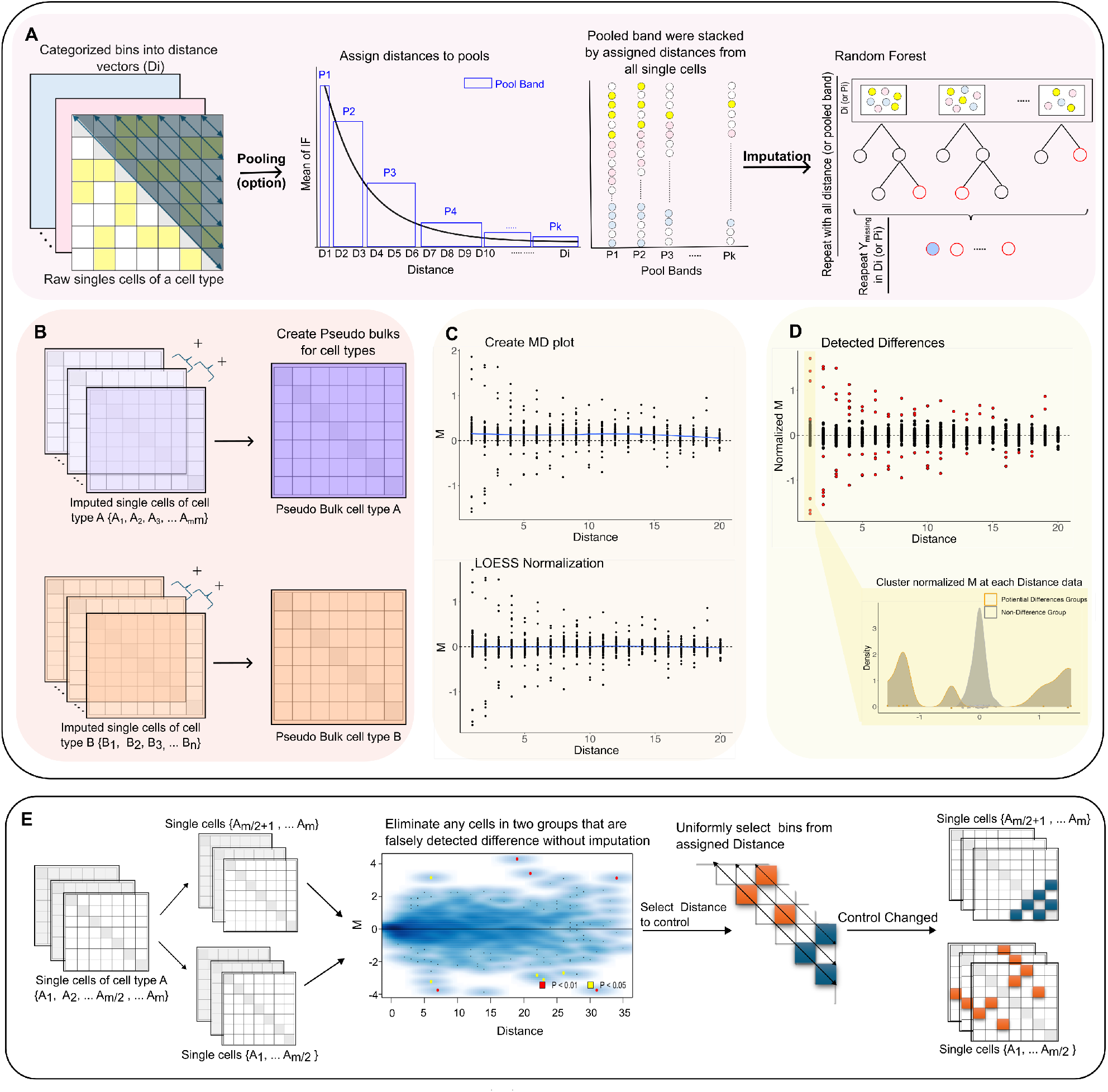
Overview of scHiCcompare. (A) Imputation. Interaction frequencies are organized into distance-based data vectors according to the physical genomic distance between two loci. With the pooling option enabled, adjacent genomic distances from all single-cell Hi-C matrices are grouped into pool bands in a progressive (or Fibonacci) style. Missing values (or zeros) in each genomic distance or pool band are imputed using the random forest method. (B) Pseudo-bulk creation. The imputed single-cell matrices of each group are summed to generate a pseudo-bulk matrix for each cell type. (C) Normalization. Two pseudo-bulk matrices are normalized using LOESS regression applied to the mean difference-distance (MD) plot. (D) Differential test. At each genomic distance, the normalized log fold changes are clustered into ‘difference’ and ‘non-difference’ groups. The ‘difference’ group (highlighted in red) is identified by filtering out the ‘non-difference’ group, which is assumed to follow a standard normal distribution. False positives are controlled by applying a log fold change threshold. (E) Defining controlled changes. Two equally sized groups of single-cell Hi-C data were randomly drawn from the same cell type-specific group. Small subsets of bin pair coordinates from each group (after eliminating unqualified bins) were selected to simulate up and down-regulated controlled changes. This results in two groups representing different conditions with a known gold standard for differential interaction.

Initially, interaction frequencies are categorized based on the physical genomic distance between two loci, generating distance data. With optional pooling techniques, this data can be aggregated into ‘pooled bands,’ capturing the distance-dependent decay of chromatin interaction frequencies. For each distance category or pooled band, missing values are imputed using random forest models (Figure 1A). Next, the imputed scHi-C matrices are summed to generate pseudo-bulk matrices for the two conditions or cell type groups (Figure 1B). These pseudo-bulk matrices are then jointly normalized using LOESS regression (Figure 1C). Inspired by our previously developed method from HiCcompare, [8], we detect differential chromatin interactions (DCI) by clustering the log fold changes (M values) across genomic distances from the mean difference-distance (MD) plot (Figure 1D). These log fold changes are classified into clusters of non-differences and significant differences using a GMM. Our results demonstrate that scHiCcompare outperforms baseline differential tests and dedicated methods on simulated controlled change data and yields biologically meaningful results on experimental data. Moreover, scHiCcompare is robust to variations in cell count and fold changes. Implemented as the R package scHiCcompare, our method enables downstream analysis and interpretation of scHi-C data.

### Imputation using Random Forest strategies improves differential analysis

A key feature that enhances our algorithm’s performance is the random forest imputation combined with pooling techniques. We aim to demonstrate how this method improves performance compared to simple mean imputation, which replaces missing interaction frequencies with the average from bins at the same distance. We use controlled changes within the scHiCcompare workflow, including normalization and differential detection, while varying the imputation methods (random forest with no pooling, progressive pooling, Fibonacci pooling, and mean distance imputation) to assess their impact across different scenarios with varying single-cell numbers across 22 chromosomes.

Before imputation, the single-cell matrix heatmaps are highly sparse, particularly at higher resolutions like 200 kb (Figure 2A). After summing to create pseudo-bulk matrices, both sparsity and noise persist, but patterns indicative of topologically associating domains (TADs) begin to emerge at both resolutions. Following imputation, matrices at 1 Mb resolution appear more complete with reduced noise, while 200 kb matrices show increased density around diagonals within a 2 Mb genomic distance range. Imputed pseudo-bulk matrices at both resolutions also reveal clearer potential TAD structures. These results highlight the effectiveness of our imputation method in reducing sparsity and recovering missing biological information in both single-cell and pseudo-bulk matrices.

**Figure 2.**
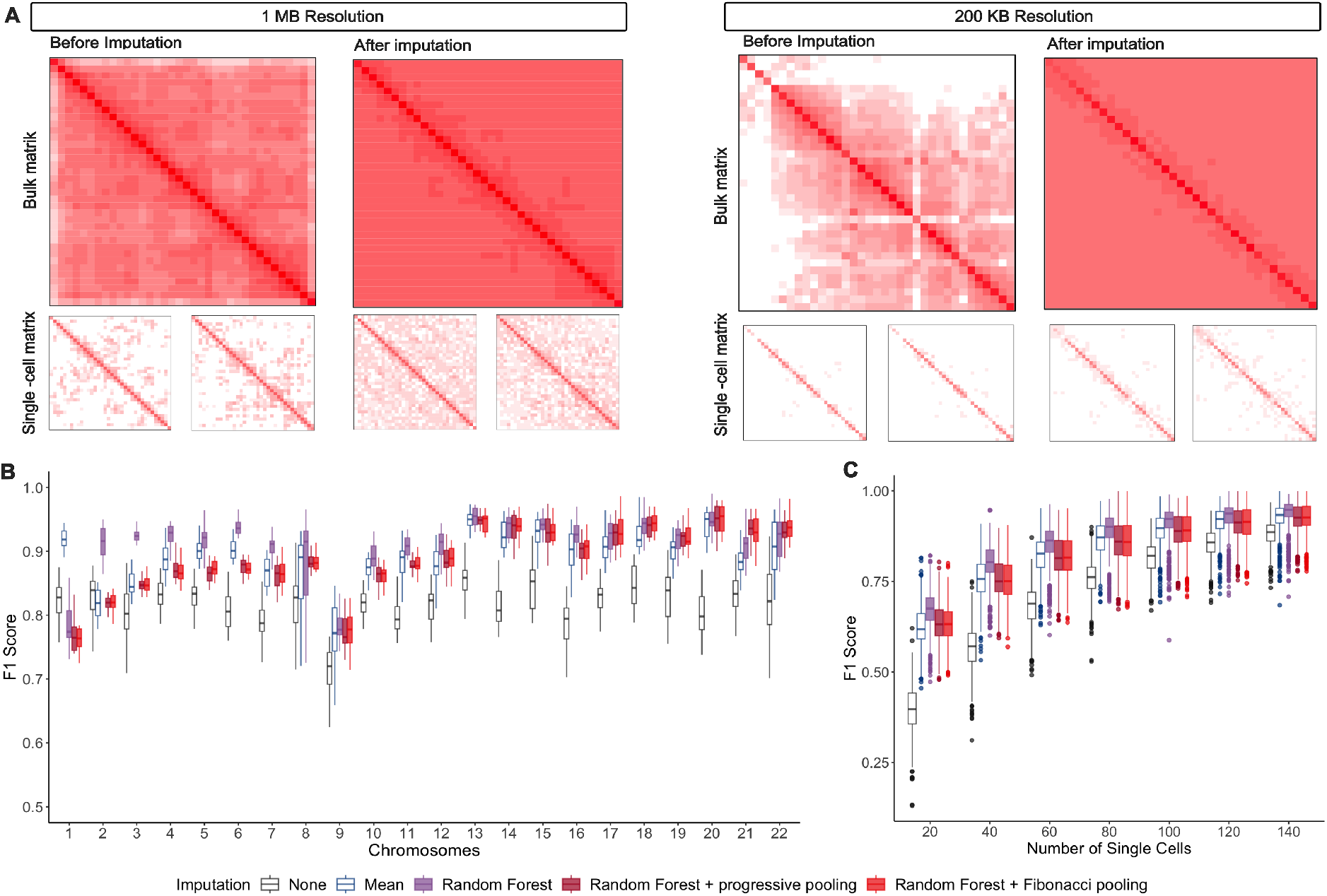
Random forest imputation with pooling technique. (A) The heatmaps of pseudo-bulk and two single-cell matrix samples are shown before and after the imputation of 100 single-cell matrices from the oligodendrocytes (ODC) cell type group, displayed at 1 Mb (from 1 to 36 Mb) and 200 kb resolutions (from 1 to 7 Mb). (B) Boxplots show the scHiCcompare workflow performance (F1 score) across each chromosome, with different imputation methods (n = 20 iterations for each chromosome). (C) Boxplots display the scHiCcompare workflow performance (F1 score) across all chromosomes when applying different imputation methods to controlled change data with various numbers of cells (20, 40, 60, 80, 100, 120, and 140 cells) in each group. The plot includes 20 iterations for each of the 22 chromosomes. The imputation methods in (B) and (C) are presented in left-to-right order: observation, mean of distance (Mean), random forest with no pooling (Random Forest), random forest with Fibonacci pooling, and random forest with progressive pooling.

Across all chromosomes, applying any imputation method generally improves workflow performance, with an average F1 score increase of 0.08 (Figure 2B). Among the methods used, the random forest imputation strategies (none, progressive, and Fibonacci pooling) consistently outperformed the baseline mean distance imputation method in most chromosomes. With the longer chromosomes (such as chromosomes 2 to 15), the random forest without pooling method exhibited the highest performance (with a mean increase of 0.097 F1 score). For the shorter chromosomes with fewer observed interaction frequencies (such as chromosomes 16 to 22), the random forest with progressive and Fibonacci pooling method achieved the highest F1 scores (with a mean increase of 0.11 F1 score) (Figure 2B). Thus, the random forest imputation, with optional pooling techniques, not only outperforms the baseline imputation method but also demonstrates robustness across chromosomes of varying lengths.

Across all chromosomes, random forest imputation without pooling consistently enhanced algorithm performance across different single-cell sample sizes (Figure 2C). For sample sizes below 80, incorporating pooling techniques improved the F1 score by at least 0.2 compared to workflows without imputation. From 80 cells onward, scHiCcompare’s performance stabilized, showing minimal changes in the median F1 score and narrower 95% confidence intervals: Random Forest - 0.91 [0.91, 0.92]; Random Forest with progressive pooling - 0.89 [0.89, 0.90]; and Random Forest with Fibonacci pooling - 0.89 [0.89, 0.90]. Additionally, when evaluating imputation methods on 100 cells at different resolutions (Supplemental Figure S2), the median of imputed interaction frequency (IF) values closely matched the original data across most genomic distances. Random forest models with progressive and Fibonacci pooling produced distributions closest to the observed IF distribution with targeted genomic distances (Supplemental Figure S3) at most resolutions. Therefore, the random forest with optional pooling techniques effectively boosts F1 scores, particularly for smaller sample sizes, while preserving the original interaction frequencies across genomic distances.

### Normalization improves differential analysis

We investigated whether alternative normalization methods could replace our LOESS normalization and enhance our algorithm. To achieve this, we compared the scHiCcompare (LOESS) normalization method with the BandNorm [17] and scHiCNorm [15] normalization methods. First, we normalized two groups of oocyte-derived cell types using the selected alternative methods. Then, we simulated controlled changes in the ODC data to create two condition groups (see Methods) on which the differential detection step was applied.

As expected, the MD plot before normalization reveals a noticeable bias at greater distances, which contradicts the expected distance effect (Figure 3A). The scHiCcompare method removes this bias most effectively, bringing the MD values closer to the baseline (M = 0) and reducing extreme log fold changes for loci at greater distances. In contrast, the other normalization methods show a deviation in the MD curve tail from the baseline (M = 0), failing to fully normalize M values at longer distances (beyond 20 Mb). Thus, scHiCcompare effectively reduces extreme values and mitigates distance-related biases between corresponding bin pairs, outperforming the other methods.

**Figure 3.**
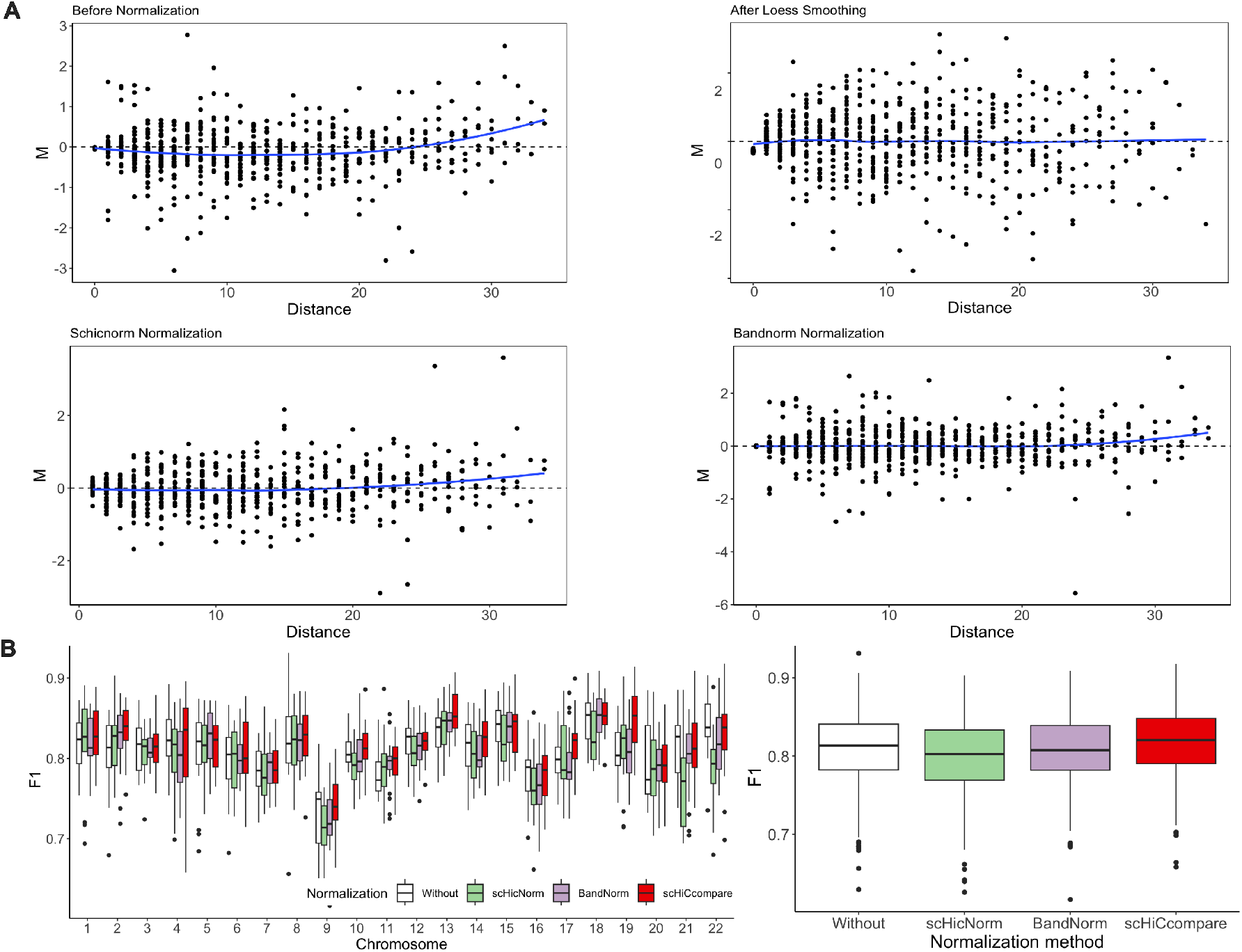
Normalization of scHi-C data. (A) MD plots visualizing the effect of different normalization methods of scHiCcompare (LOESS), scHiCNorm, and BandNorm) versus before any normalization applied from controlled changed ODC datasets. (B) The left figure compares the performance of scHiCcompare (LOESS), scHiCNorm, and BandNorm in terms of F1 score across each of the 22 chromosomes, both before and after normalization. The right figure presents F1 score comparisons across all chromosomes. The performance of all methods for each chromosome was measured over 20 iterations.

When comparing all normalization methods across individual chromosomes (Figure 3B), scHiCcompare achieved the highest performance in 12 chromosomes, BandNorm [17] in 3 chromosomes, and scHiCNorm [15] in 3 chromosomes. This indicates that scHiCcompare normalization consistently enables the differential test to achieve the highest F1 scores across most chromosomes. For all chromosomes combined (Figure 3B), the workflow using scHiCcompare also produces the highest median F1 score (0.820 [range: 0.66-0.92]), along with superior precision, sensitivity, specificity, and MCC scores (Supplemental Figure S4). Therefore, scHiCcompare normalization using LOESS regression proves to be the most effective and consistent method overall.

### Differential analysis of pseudo-bulk scHi-C data outperforms baseline methods

To evaluate the performance of our pseudo-bulk differential test, we compared it with other parametric (T-test, Negative Binomial test) and non-parametric (Wilcoxon rank-sum and Kolmogorov-Smirnov test) methods. This comparison was conducted using simulated difference data that included controlled changes with varying fold changes across different genomic distances spanning all chromosomes. In each case, we measured the performance of these differential detection tests on IFs within distances below 20 Mb (see Methods). All tests were followed by a workflow of normalization and differential testing. While scHiCcompare normalizes and analyzes pseudo-bulk matrices, parametric and non-parametric tests compare IF vectors from bin pairs across scHi-C matrices, requiring joint normalization across multiple matrices. We used LOESS-based multiHiCcompare [9] for joint normalization before non-parametric and parametric tests.

When comparing chromatin interaction differences at varying fold changes (FC) (Figure 4A), the scHiCcompare differential method consistently achieved the highest median F1 scores across all chromosomes (FC = 3: 0.82; FC = 6: 0.87; FC = 9: 0.87). In terms of precision, sensitivity, and specificity metrics, scHiCcompare consistently ranked in the top two across all chromosomes (Supplemental Figure S5). Although the Kolmogorov-Smirnov (KS) test performed well in higher fold change scenarios (FC = 6 and FC = 9), it tended to detect more false positives with lower sensitivity in cases of differences with a small fold change (Supplemental Figure S5). Conversely, the Wilcoxon rank-sum test showed the lowest F1 scores across all fold changes, while the T-test and Negative Binomial models performed similarly with only moderate improvements as the fold change increased. Therefore, scHiCcompare’s differential method is the most consistent and reliable option. It accurately detects differential chromatin interactions (DCIs) and controls false positives across various fold changes and chromosomes, outperforming both parametric and non-parametric tests.

**Figure 4.**
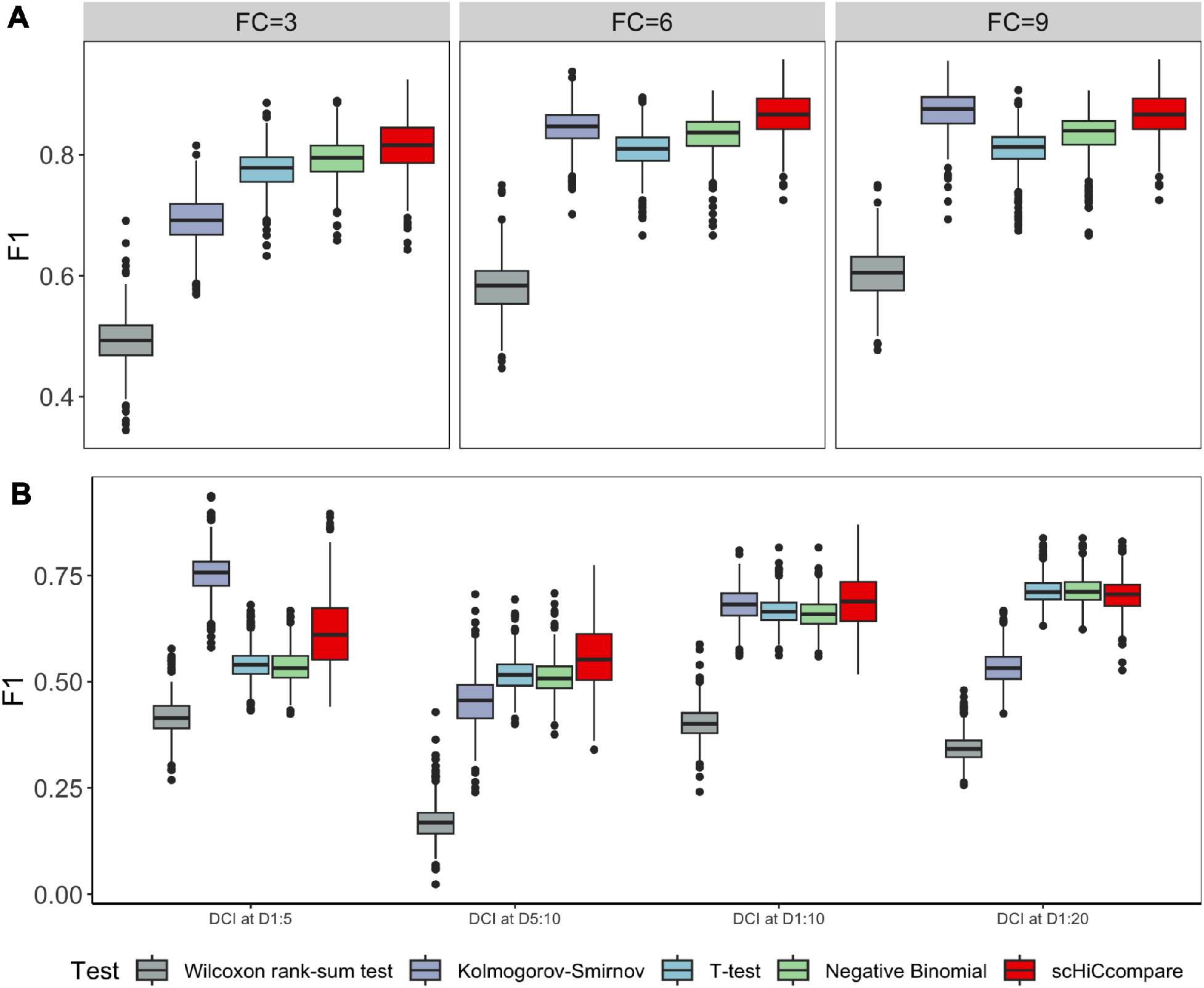
Compare scHiCcompare differential test method with baseline methods. (A) These boxplots show F1 scores comparing the scHiCcompare pseudo-bulk differential methods with baseline methods (Wilcoxon rank-sum test, Kolmogorov-Smirnov test, T-test, and Negative Binomial test) at different fold changes (FC = 3, FC = 6, FC = 9) across all chromosomes. (B) These boxplots show the overall performance of the differential methods of scHiCcompare compared with baseline methods in various scenarios of control-changed DCIs simulated at different genomic distance ranges, including 1 Mb to 5 Mb (DCI at D1:5), 5 Mb to 10 Mb (DCI at D5:10), 1 Mb to 10 Mb (DCI at D1:10), and 1 Mb to 20 Mb (DCI at D1:20), across all chromosomes. For all boxplots in (A) and (B), each chromosome’s reported F1 score includes 20 iterations.

When comparing cases of controlled chromatin interaction differences across various distance ranges (fold change = 3) in all chromosomes, the scHiCcompare pseudo-bulk differential method demonstrates superior average F1 scores with narrower 95% confidence intervals in most cases (DCI at distance 1-5 Mb: mean = 0.62 [0.61, 0.62]; DCI at distance 5-10 Mb: mean = 0.56 [0.55, 0.57]; DCI at distance 1-10 Mb: mean = 0.69 [0.68, 0.70]; DCI at distance 1-20 Mb: mean = 0.70 [0.70, 0.71]) (Figure 4B). The Kolmogorov-Smirnov test performs second-best, showing the highest performance when differences occurred at distances of 1-5 Mb, but its effectiveness was inconsistent and declined in other scenarios. Methods like the Wilcoxon test performed the worst in all scenarios. The T-test and Negative Binomial tests exhibited similar performance, which improved with wider genomic distance ranges of controlled changes, but generally remained below that of scHiCcompare (Figure 4B). This highlights the robustness of the scHiCcompare pseudo-bulk differential method in accurately detecting chromatin interaction differences across various genomic distances compared to other methods.

### scHiCcompare outperforms Snap-HiC-D and scHiCDiff methods for differential scHi-C analysis

We compared our scHiCcompare workflow with existing scHi-C differential analysis methods, SnapHiC-D [11] and scHiCDiff [12]. We applied all three algorithms to a control changes dataset on chromosome 22 with 20 iterations (see Methods). Due to limitations with the resolutions supported by each method, we compared scHiCcompare (using progressive pooling with random forest imputation) with SnapHiC-D at a 10 kb resolution, as SnapHiC-D operates only at 10 kb. scHiCDiff demands substantial runtime and memory at 10 kb resolutions, so we performed comparisons at the lower resolutions of 200 kb, 500 kb, and 1 Mb on chromosome 22.

After 20 iterations, scHiCcompare achieved an average F1 score at least twice that of SnapHiC-D and scHiCDiff (Figure 5A) across various fold changes and resolutions. SnapHiC-D and scHiCDiff showed low sensitivity, indicating a high rate of false positives in most cases (Supplemental Figure S6). In contrast, scHiCcompare demonstrated robust overall performance, attaining a mean F1 score of at least 0.6 across most resolutions, with the exception of the 10 kb dataset. Our method’s F1 scores reached as high as 0.9 at a 1 Mb resolution across all fold changes, a 500 kb resolution with fold changes of 6 and 9, and a 200 kb resolution with a fold change of 9. Overall, scHiCcompare demonstrated improved performance as both resolution and fold change increased, outperforming the two existing scHi-C differential analysis methods.

**Figure 5.**
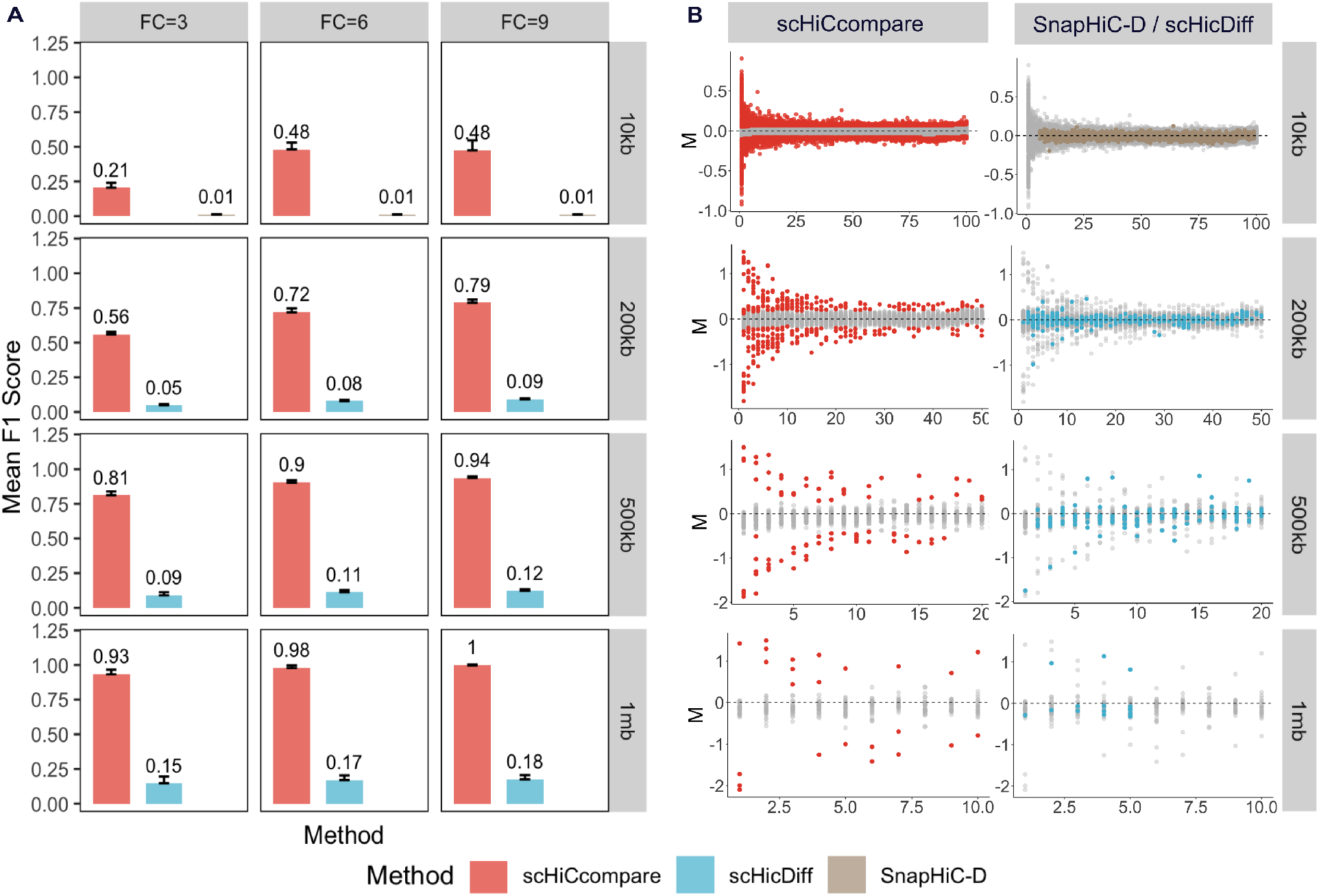
scHiCcompare compared with SnapHiC-D and scHiCDiff methods. (A) The bar plots show the mean F1 scores for chromosome 22 (20 iterations) in detecting differential chromatin interactions (DCIs) in controlled difference data at various fold changes (FC = 3, 6, 9) and resolutions (10 kb, 200 kb, 500 kb, and 1 Mb). Error bars represent the standard deviations of F1 scores across multiple runs. (B) MD plots visualize the differential results for a sample run of ODC data with simulated differences at fold change 3 across four resolution scales: 10 kb, 200 kb, 500 kb, and 1 Mb. The left column displays results from scHiCcompare, while the right column presents results from SnapHiC-D and scHiCDiff. Colored points indicate detected DCIs for each method (red for scHiCcompare, blue for scHiCDiff, and brown for SnapHiC-D), while grey points represent ‘non-difference’ interactions.

At all resolutions, scHiCcompare consistently captured differential chromatin interactions with pseudo-bulk’s log fold changes significantly deviating from the baseline (M = 0) (Figure 5B). In contrast, SnapHiC-D and scHiCDiff tended to capture small log fold changes, the majority of which center around the baseline (M = 0). These MD plots provide further evidence that scHiCcompare effectively identifies significant DCIs with high log fold changes, while the other methods detect more false positives with lower log fold changes.

### scHiCcompare provides biological insights

To further evaluate the biological relevance of the differential chromatin interactions (DCIs) identified by scHiCcompare, we examined whether these DCIs provide biological insights in experimental data. Since scHiCcompare has shown to be robust across groups with varying numbers of cells (Supplemental Note S4), we applied the scHiCcompare workflow to detect DCIs in human brain cells [18] and mouse cells [19], both of which include different numbers of cells for each cell type group.

To see if scHiCcompare can catch the unique biology of excitatory cells, we conducted multiple comparisons between pairs of inhibitory neuron cells (Ndnf and Vip) and excitatory neuron cells (L2/3, L4, L5, and L6) at 1 Mb resolutions using the no pooling random forest imputation setting (outlier significance level = 0.06, other parameters set to default; see Methods). Our analysis revealed cell type-specific chromatin interactions consistent with previous findings, where chromatin loops between the SATB2 promoter region and the adjacent LINC01923 locus are only found in excitatory cells, but not inhibitory neuron types [18]. In particular, we identified significant DCI with two boundary regions overlapping with the genomic loci of SATB2 and LINC01923 in most of our comparisons between inhibitory and excitatory neurons. The majority of these DCIs exhibited adjusted fold changes consistent with higher normalized interaction frequencies in excitatory neurons compared to inhibitory neurons (Supplemental Table S1).

Next, to gain biological insights and explore relevant epigenetic features specific to cell types, we identified chromatin differences between astrocytes (Astro) and endothelial cells (Endo) on chromosome 22 at a 50 kb resolution [18] using the scHiCcompare workflow with progressive pooling imputation. We identified 9,363 DCIs at this resolution covering regions that overlap with previously reported differential chromatin domain boundaries in the human prefrontal cortex, especially in comparisons between Astro and Endo cells [18] (Supplemental Table S2). Additionally, we compared the histone marker scores (H3K4me3, H3K4ac, and CTCF) of Astro and Endo cells in regions with up-regulated and down-regulated differential boundaries. Although not all score comparisons were statistically significant, cells with higher interaction frequencies in DCI regions generally showed increased levels of histone and CTCF signals in those regions (Supplemental Table S3). (Supplemental Table S3). In an example of a DCI region with higher interaction frequencies in Astro (Supplemental Figure S7), we observed consistent patterns in histone modification data: H3K4me3 peaks in Astro were higher than in Endo cells (highlighted in purple in Supplemental Figure S6), and Astro boundaries also displayed elevated H3K4ac signals compared to endothelial cells. These findings suggest that cells with higher interaction frequencies in DCIs often have increased levels of specific histone modification signals relative to their counterparts.

To extend the evaluation of our method beyond human data and assess its biological relevance, we analyzed chromatin interactions between active immature oocytes (non-surrounded nucleolus, NSN) and mature oocytes (surrounded nucleolus, SN) in mice, spanning 19 chromosomes at a 200 kb resolution [19] using the scHiCcompare workflow with progressive pooling random forest imputation. Our analysis identified 148,888 DCIs across all chromosomes, covering distances from 400 kb to 10 Mb. The normalized fold change magnitude of these DCIs, both up-regulated and down-regulated fold change, consistently exhibited a mean of around 1.2 across all genomic distances, suggesting relatively small differences in IF between SN and NSN oocytes at all distances analyzed (Supplemental Table S4). Notably, in all 19 chromosomes, the detected differences revealed more positive log fold changes, indicating a higher IF in SN compared to NSN oocytes for interactions above the 400 kb range (Supplemental Table S5). These findings are consistent with the original report, which noted that while the overall contact probability (determined by genomic distance) is similar between immature and mature oocytes, mature oocytes exhibit a higher number of long-range (>400 kb) contacts [19].

## Discussion

We developed scHiCcompare for the differential analysis of single-cell Hi-C data. scHiCcompare imputes single-cell matrices while maintaining both genomic distance effects and cell-specific variability, employing a pseudo-bulk strategy to normalize pseudo-bulk matrices from two groups of scHi-C data. The subsequent differential detection method outperforms baseline comparison methods (parametric and non-parametric differential tests), as well as SnapHiC-D [11] and scHiCDiff [12], in detecting true DCIs while effectively controlling for false positives. scHiCcompare demonstrates robust performance across various genomic distance ranges, fold changes, data resolutions, and single-cell counts in each group.

In addition to its statistical advantages, scHiCcompare addresses limitations of existing methods. scHiCcompare is compatible with data at various resolutions, whereas SnapHiC-D can only analyze 10 kb scHi-C data. Additionally, the imputation, normalization, and differential detection steps in SnapHiC-D require separate scripts, making its workflow challenging to execute from start to finish. While scHiCDiff supports multiple resolutions, it becomes prohibitively slow at higher resolutions, such as 10 kb (requiring several days). scHiCcompare workflow with pooling strategies run more faster than scHiCDiff and SnapHiC-D in various resolutions (Supplemental Table S6).

The performance of our imputation method can be influenced by the percentage of missing values (zeros) and the availability of observed interaction frequencies at each genomic distance. To accommodate these factors, we offer different imputation approaches (with and without pooling). Random forest imputation without pooling is effective when there are sufficient existing IF observations (e.g., at least 1,500 IF values at each target genomic distance) and when the missing value percentage is relatively low (e.g., no more than 80%) at each distance. This approach is particularly efficient for longer chromosomes (such as chromosomes 1 to 15) at lower resolutions (above 500 kb). In more extreme cases, pooling techniques can gather additional observations across pooled bands, enhancing imputation but with trade-offs in runtime and memory usage.

While our imputation method preserves the distance effect and correlation between cells, it does not consider spatial relationships between loci, assuming that missing values are random. Imputation reliability could be further improved by incorporating spatial localization, such as k-nearest neighbor imputation. Additionally, our study focuses on differences with fold changes of 3, 6, and 9, following the scHiCDiff study, which SnapHiC-D does not examine. Consequently, our method may overlook less pronounced changes with smaller fold changes. Future improvements could involve adaptively optimizing the filtered log fold change threshold to better capture low log fold changes at each genomic distance.

## Methods

### Overview

The scHiCcompare workflow consists of four main steps: (1) scHi-C data imputation using random forest applied to genomic distances or pooled bands, (2) generation of pseudo-bulk matrices by summing the interaction frequencies (IF) of cells for each cell type group, (3) joint LOESS normalization of the two pseudo-bulk matrices, and (4) detection of differential chromatin interactions.

### Input

The scHi-C data should be pre-clustered to define two groups of chromosome-specific cells (or conditions) for comparison. The data format is a modified sparse upper triangular matrix with five columns: the chromosome of the first interacting region, (2) the starting coordinate of the first region, (3) the chromosome of the second interacting region, (4) the starting coordinate of the second region, and (5) the interaction frequency between the interacting regions. Each cell’s scHi-C data for a given condition should be saved in a separate text file, with files for each group stored in distinct folders to serve as inputs for scHiCcompare.

### Imputation on bands

To impute missing scHiC data, random forest models are employed. The imputation step is applied independently to single-cell groups from each cell type. Initially, we categorize interaction frequencies into distance-specific data, based on the physical genomic distance between two loci (off-diagonal vectors on Hi-C maps). Imputation is then performed on each genomic distance data individually or pooled into bands combining similar interaction frequencies.

Two versions of pooling techniques are implemented to gather genomic distances across all group-specific single-cell matrices into consecutive pooled bands. Pooling addresses the genomic distance effect, enabling interaction frequencies with similar distance effects to be grouped for analysis. We implemented two pooling techniques: progressive and Fibonacci pooling (Figure 1A).

1. In progressive pooling, subsequent pool bands contain interaction frequencies from larger genomic distance ranges, with the distance range increasing linearly in each subsequent pool. For example, in units of resolution, the first pool *P*_1_ contains first genomic distance (*D*_1_); the second pool *P*_2_ contains second and third off-diagonal interaction frequencies (*D*_2_) and (*D*_3_); the third pool *P*_3_ contains fourth, fifth, and sixth genomic distance (*D*_4_), (*D*_5_), (*D*_6_), etc.,
2. In Fibonacci pooling, the distance range increases according to the Fibonacci sequence (Supplemental Methods).

We assume that missing data in the single-cell contact matrix is missing at random and can thus be imputed using the remaining observed data. To address this, random forest imputation is adapted for each genomic distance or pooled band to infer missing values. The random forest model is trained on genomic-or band-specific existing contact data, excluding extreme values outside the interquartile range (IQR), with cell identity (indicating the specific single cell in which the contact occurs) included as a feature. Predictions are then applied across all interaction frequencies within the distance or pooled band to impute missing values (Supplemental Methods). This approach ensures that imputed frequencies reflect the genomic distance effect while preserving correlations across single cells within each cell type.

### Pseudo-bulk

Imputed single-cell matrices within each group are summed to generate two pseudo-bulk matrices for each cell type. Assuming that the interaction frequency between two loci in a single cell has a similar distribution across all cell type or condition-specific scHi-C matrices, summing these interactions gathers the maximum amount of information.

### Normalization

Like most Hi-C techniques, single-cell Hi-C data has global and local biases in chromatin interaction due to technology imperfections and DNA sequence properties. To mitigate this effect, we apply the HiCcompare normalization method to normalize the two imputed pseudo-bulk matrices jointly. Briefly, the data is represented on a mean-difference (MD) plot, based on the idea of the MA plot (Bland-Altman plot) [20] used for visualization of gene expression differences. The **M**ean log difference between the two datasets (*M*) is calculated as *M* = *log*_2_(*IF*_2_*/IF*_1_), where *IF*_1_ and *IF*_2_ represent interaction frequencies of the first and second Hi-C datasets, respectively. Additionally, *D* represents the **D**istance between two interacting regions in terms of the resolution of the scHi-C data. To address global biases, a LOESS regression curve is fitted through the MD plot and used to adjust the *M* differences around a baseline of *M* = 0 [8].

### Differential Detection

This step involves clustering the normalized log fold changes of interaction frequencies at each genomic distance into ‘difference’ and ‘non-difference’ clusters. At each genomic distance, we assume that, in the absence of significant DCI between two groups (non-difference clusters), the normalized log fold changes follow a standard normal distribution centered around 0 [8]. To test this assumption, we first conduct a Shapiro-Wilk test to determine whether the M values at a given genomic distance follow a standard normal distribution.

If the M values at a distance are found not to follow a normal distribution, this indicates the presence of ‘difference’ DCIs that form separate distributions deviating from the ‘non-difference’ normal distribution. In this case, these differences are detected by applying Gaussian Mixture Model (GMM) clustering, which classifies the M values into two groups: (1) the normal distribution group representing non-significantly different M values, and (2) the outlier distributions, representing up and down-regulated M values, which indicate potential significant differences (Supplemental Methods). We further filter these differences using a predefined log fold change threshold to exclude M values with low log fold changes.

If the Shapiro-Wilk test confirms that the M values follow a normal distribution, it suggests there are no significant differences or only a few differences insufficient to form distinct distributions. In this case, these few differences are assumed to be outliers of the normal distribution, which are identified by the differential analysis in the HiCcompare method [8]. Briefly, the distance-specific M values (after filtering out low-average expression interaction frequencies) are transformed into Z-scores and corresponding p-values using the standard normal distribution. This method effectively identifies significant outlier differences in values between two cell type groups.

In the default settings for experimental studies, the imputation step is set to perform 10 multiple imputations using the selected pooling style. The log fold change threshold for identifying differential clusters is set to 0.8 (equivalent to approximately a 2-fold change), and the significance level for outlier detection is set to 0.5.

## Controlled changes

### Controlled Changes Definition in Raw Data

To evaluate the performance of differential interaction detection methods, controlled interaction differences are introduced. Assuming minimal to no differential interactions among cells of the same cell type, two groups of scHi-C matrices are created by randomly drawing data from the same cell type. Controlled differences are then introduced in the two groups as follows (Figure 1E):

1. Apply the HiCcompare differential method [8] to identify M value outliers, or false positive difference bin pairs, between un-imputed and normalized scHi-C pseudo-bulk matrices from the two initial groups of the same cell type.
2. Select a genomic distance range to introduce controlled changes. By default, this range is set from 1 to 10 Mb, excluding the bin pairs at the diagonal.
3. Identify qualified bins for controlled changes by filtering out unqualified bins, including:

- Bins identified as false positives in step 1
- Bins with low average expression, where the average normalized interaction frequency, 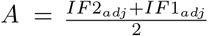, is below the 10th percentile of all bin pairs’ A values
- Bins with large differences between pairs, where the *M* value, 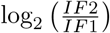, is greater than 1 or less than -1

4 Uniformly select a number of bin pairs for controlled changes from the qualified bins identified in step 3. By default, the number of bin pairs selected is 10% of the qualified bins.

The selected bins are randomly divided into two equal sets: up and down-regulated (in group 2 versus group 1 comparison). For up-regulated bins (positive log fold change), the interaction frequencies in group 2 are multiplied by a fold change, while those in group 1 remain unchanged. The opposite was done for down-regulated bins. These simulated controlled changes are used as a “gold standard” for benchmarking the performance of differential interaction detection methods.

In the default setting, differences are simulated to create two benchmark groups, each comprising 100 cells, using oocyte-derived cell type scHi-C data at a 1 Mb resolution. Due to the extreme sparsity at higher genomic distances, the scHiCcompare workflow focuses on analyzing scHi-C data within the 1 to 10 Mb genomic distance range. Controlled changes are defined within this range, with a fold change of 3. One of four factors is then varied—fold change, resolution, sample size, or controlled DCI’s genomic distance ranges—for targeted comparisons. For each simulation setting, 20 iterations per chromosome are generated to evaluate performance.

## Data sources

To evaluate the performance of the scHiCcompare workflow, we utilized published datasets, including human brain prefrontal cortex cells [18] and mouse oocyte-to-zygote cells [19]. Lee et al. scHi-C data included 14 human brain prefrontal cortex cell types: Astro, Endo, L2/3, L4, L5, L6, MG, MP, Ndnf, ODC, OPC, Pvalb, Sst, and Vip, originating from two donors aged 21 and 29 years, across a total of five sequencing libraries. Data at a 10 kb resolution were downloaded from https://salkinstitute.app.box.com/s/fp63a4j36m5k255dhje3zcj5kfuzkyj1/folder/82405563291. Data at a 1 Mb resolution were accessed using the download_schic() function of Bandnorm [17] in R. The Flyamer et al. dataset includes mouse cells in two conditions: oocyte and zygote. We downloaded the dataset from GEO [21] with data accession GSE80006 at 40 kb and 200 kb resolutions.

## Code availability

scHiCcompare R package is available on GitHub at https://github.com/dozmorovlab/scHiCcompare.

## References

[1] A.D. Schmitt, M. Hu, B. Ren, Genome-wide mapping and analysis of chromosome architecture, Nature Reviews Molecular Cell Biology 17 (2016) 743–755.

[2] B. Bonev, G. Cavalli, Organization and function of the 3D genome, Nature Reviews Genetics 17 (2016) 661–678.

[3] Y. Li, M. Hu, Y. Shen, Gene regulation in the 3D genome, Human Molecular Genetics 27 (2018) R228–R233.

[4] P.H.L. Krijger, W. De Laat, Regulation of disease-associated gene expression in the 3D genome, Nature Reviews Molecular Cell Biology 17 (2016) 771–782.

[5] E. Lieberman-Aiden, N.L. Van Berkum, L. Williams, M. Imakaev, T. Ragoczy, A. Telling, I. Amit, B.R. Lajoie, P.J. Sabo, M.O. Dorschner, others, Comprehensive mapping of long-range interactions reveals folding principles of the human genome, Science 326 (2009) 289–293.

[6] T. Nagano, Y. Lubling, T.J. Stevens, S. Schoenfelder, E. Yaffe, W. Dean, E.D. Laue, A. Tanay, P. Fraser, Single-cell hi-c reveals cell-to-cell variability in chromosome structure, Nature 502 (2013) 59–64.

[7] T. Nagano, Y. Lubling, C. Várnai, C. Dudley, W. Leung, Y. Baran, N. Mendelson Cohen, S. Wingett, P. Fraser, A. Tanay, Cell-cycle dynamics of chromosomal organization at single-cell resolution, Nature 547 (2017) 61–67.

[8] J.C. Stansfield, K.G. Cresswell, V.I. Vladimirov, M.G. Dozmorov, HiCcompare: An r-package for joint normalization and comparison of HI-c datasets, BMC Bioinformatics 19 (2018) 1–10.

[9] J.C. Stansfield, K.G. Cresswell, M.G. Dozmorov, multiHiCcompare: Joint normalization and comparative analysis of complex hi-c experiments, Bioinformatics 35 (2019) 2916–2923.

[10] M.N. Djekidel, Y. Chen, M.Q. Zhang, FIND: difFerential chromatin INteractions detection using a spatial poisson process, Genome Research 28 (2018) 412–422.

[11] L. Lee, M. Yu, X. Li, C. Zhu, Y. Zhang, H. Yu, Z. Chen, S. Mishra, B. Ren, Y. Li, others, SnapHiC-d: A computational pipeline to identify differential chromatin contacts from single-cell hi-c data, Briefings in Bioinformatics 24 (2023) bbad315.

[12] H. Liu, W. Ma, scHiCDiff: Detecting differential chromatin interactions in single-cell hi-c data, Bioinformatics 39 (2023) btad625.

[13] R. Zhang, T. Zhou, J. Ma, Multiscale and integrative single-cell hi-c analysis with higashi, Nature Biotechnology 40 (2022) 254–261.

[14] J. Zhou, J. Ma, Y. Chen, C. Cheng, B. Bao, J. Peng, T.J. Sejnowski, J.R. Dixon, J.R. Ecker, Robust single-cell hi-c clustering by convolution-and random-walk–based imputation, Proceedings of the National Academy of Sciences 116 (2019) 14011–14018.

[15] T. Liu, Z. Wang, scHiCNorm: A software package to eliminate systematic biases in single-cell hi-c data, Bioinformatics 34 (2018) 1046–1047.

[16] W.G. Jacoby, Loess:: A nonparametric, graphical tool for depicting relationships between variables, Electoral Studies 19 (2000) 577–613.

[17] Y. Zheng, S. Shen, S. Keleş, Normalization and de-noising of single-cell hi-c data with BandNorm and scVI-3D, Genome Biology 23 (2022) 222.

[18] D.-S. Lee, C. Luo, J. Zhou, S. Chandran, A. Rivkin, A. Bartlett, J.R. Nery, C. Fitzpatrick, C. O’Connor, J.R. Dixon, others, Simultaneous profiling of 3D genome structure and DNA methylation in single human cells, Nature Methods 16 (2019) 999–1006.

[19] I.M. Flyamer, J. Gassler, M. Imakaev, H.B. Brandão, S.V. Ulianov, N. Abdennur, S.V. Razin, L.A. Mirny, K. Tachibana-Konwalski, Single-nucleus hi-c reveals unique chromatin reorganization at oocyte-to-zygote transition, Nature 544 (2017) 110–114.

[20] S. Dudoit, Y. Yang, M. Callow, T. Speed, Statistical methods for identifying genes with differential expression in replicated cDNA microarray experiments, Stat. Sin (2002).

[21] R. Edgar, M. Domrachev, A.E. Lash, Gene expression omnibus: NCBI gene expression and hybridization array data repository, Nucleic Acids Research 30 (2002) 207–210.

