## Supplemental for "scHiCcompare: an R package for differential analysis of single-cell Hi-C data"

### Supplemental Methods

#### Supplemental Note S1. Pooling technique in imputation

Before imputation, the data of the single-cell matrix bin is assigned into different pooling bands in distance units. We have two main pooling designs:

1. Progressive pooling design
2. Fibonacci pooling design

##### 1. Progressive pooling

Subsequent pools contain single or strings of consecutive distances with the number of elements in the pool matching its index. Thus, the $i$-th pool band ($P_i$) includes $i$ consecutive distance elements. As the pool band index increases by 1 unit, the number of elements in the pool increases by 1 distance unit until all unit distances belong to one of $K$ pools. In the case where the last pool band ($P_K$) reaches the last distance ($D_N$) but does not contain enough distance elements, the extra distances are drawn back to the previous distances until the pool is filled. For example, if the last pool is $P_4$ and needs 4 distance elements but the last distance is $D_8$, then

$$P_1: \{D_1\}$$

$$P_2: \{D_2, D_3\}$$

$$P_3: \{D_4, D_5, D_6\}$$

$$P_4: \{D_7, D_8, D_7, D_6\}$$

If arg max$_{j \in P_{K-1}} D_j + K = N; j = \{1,2,\dots,N\}$:

$$P_i = \{D_j \mid L_{i-1} + 1 \le j \le L_{i-1} + i\}, \quad i = \{1,2,\dots,K\}$$

If arg max$_{j \in P_{K-1}} D_j + K > N; j = \{1,2,\dots,N\}$:

$$P_i = \begin{cases} \{D_j \mid L_{i-1} + 1 \le j \le L_{i-1} + i\}, & \text{if } i = 1,2,\dots,K-1 \\ \{D_j \mid K - i + 1 \le j \le K\}, & \text{if } i = K \end{cases}$$

Where:

- $P_i$ = pooling band $i$
- $D_j$ = IF values of bins in distance $j$
- $L_{i-1}$ = largest index $j$ of distance belongs to $P_{i-1}$
- $K$ = the largest index of the pool band
- $N$ = the maximum index of distance

##### 2. Fibonacci pooling

Given the Fibonacci sequence is defined as:

$$F_0 = 0; \quad F_1 = 1; \quad F_l = F_{l-1} + F_{l-2}, \text{ for } l > 2$$

The Fibonacci pooling sample contain index follow by Fibonancci sequence above. In the case where the last pool band ($P_K$) reaches the last distance ($D_N$) but does not contain enough distance elements, the extra distances are drawn back to the previous distances until the pool is filled.

If arg max$_{j \in P_{K-1}} D_j + F_K = N; j = \{1,2,\dots,N\}$:

$$P_i = \begin{cases} D_i, & \text{if } i = 0,1 \\ \{D_j \mid L_{i-1} + 1 \le j \le L_{i-1} + F_i\}, & \text{if } i = 2,3,\dots,K \end{cases}$$

If arg $\max_{j \in P_{K-1}} D_j + F_K > N$; $j = \{1,2,\dots,N\}$:

$$P_i = \begin{cases} D_i, & \text{if } i = 0,1 \\ \{D_j \mid L_{i-1} + 1 \le j \le L_{i-1} + F_i\}, & \text{if } i = 2,3,\dots,K-1 \\ \{D_j \mid N - F_K + 1 \le j \le K\}, & \text{if } i = K \end{cases}$$

**Supplemental Note S2. Random forest imputation**

Assuming that the sparsity in the single-cell matrices is due to missing values occurring at random, we adapt MICE random forest imputation to infer these missing values [1]. All values of 0 in a matrix are considered missing and are therefore subject to imputation. Given $n$ single-cell matrices of size $m \times m$ with $k$ bins assigned to the $i^{th}$ pooling band $P_i$, the imputation model is applied to a training matrix of each pooling band, $X_{P_i}$:

$$X_{P_i} = [x_{ki} \quad c_m]$$

where $x_{ki}$ represents the interaction frequency value of bin $k$ in pooling band $P_i$, located in a single cell $C_m$. Each $X_{P_i}$ contains observed values $X_{P_i}^{obs}$ and missing values $X_{P_i}^{mis}$. The imputation for each pool is performed in the following steps:

1. Remove any extreme values in $X_{P_i}$ that are less than or greater than a specified multiple of the interquartile range $(IQR = Q3 - Q1)$.
2. Fill initial values for $X_{P_i}^{mis}$ by randomly drawing from $X_{P_i}^{obs}$, creating $X_{\dot{P}_i}$.
3. Repeat this step $l$ times (number of iterations):
    a) Draw $k$ (n_tree) bootstrap samples from $X_{\dot{P}_i}$.
    b) Fit a tree to each bootstrap, resulting in $k$ (n_tree) trees. Each tree has multiple leaves, where each leaf contains some $X_{P_i}^{obs}$ values, referred to as donors.
    c) Predict the model from step b on $X_{P_i}^{obs}$ and identify which $X_{P_i}^{obs}$ values are contained in each leaf, serving as donors within each leaf.
    d) Predict the model from step b on $X_{P_i}^{mis}$ and determine the leaves to which $X_{P_i}^{mis}$ values belong in $k$ (n_tree) trees.
    e) Impute each $X_{P_i}^{mis}$ by randomly selecting an $X_{P_i}^{obs}$ value within the same $k$ leaves as identified in step d.
    f) Replace the original missing values $X_{P_i}^{mis}$ in $X_{\dot{P}_i}$ with the imputed values from step e.
4. Repeat steps 1–3 for $m$ times, resulting in $m$ imputed sets.

If the existing observations within each pool exhibit collinearity or consist of a single constant value, the $X_{P_i}$ band will be imputed by its mean. Our imputation method primarily targets a specific range of genomic distances; by default, this range is set to 1–10 MB, within which full imputation is performed. For distances outside this primary range, if the percentage of missing values exceeds a predefined threshold (set at 95% by default), imputation will not be performed by the random forest model but will instead be completed using the mean value.

**Supplemental Note S3. Gaussian Mixture Model (GMM) cluster differential test**

We assume that if there are no significant differences between bin pairs of two different condition groups (non-difference clusters), their normalized log fold change (M values) is assumed to follow a standard normal distribution centered around 0. The 'difference' cluster consists of M values that deviate from the 'non-difference' cluster's standard normal distribution. Therefore, the differential bins can be identified by filtering out this 'non-difference' cluster classified by the Gaussian Mixture Model (GMM) method at each genomic distance. The GMM is set to classify the data into three clusters ($j = 3$):

1. The middle cluster (with the second ranked estimated mean that has closest mean to 0) represents the non-difference group.

2. The other two clusters correspond to up-regulated and down-regulated M value distributions, representing differences.

Given that we have $D_1, D_2, \dots, D_j$ genomic distances, each containing an interaction frequency of $x_{1j}, x_{2j}, \dots, x_{ij}$, $n = 1,2,\dots,n$ as random variables drawn i.i.d., the genomic distance $D_j$ is assumed to follow the density:

$$P_{D_j}(x) = \sum_{k=1}^{3} \pi_k \, \phi(x; \mu_j k \Sigma_k)$$

Each cluster component ($k = 3$) is assumed to have a multivariate Gaussian distribution, where:

- $\phi(x; \mu_k, \Sigma_k) = \frac{1}{(2\pi|\Sigma_k|)^{1/2}} \exp\left(-\frac{1}{2}(x-\mu_k)^T \Sigma_k^{-1}(x-\mu_k)\right)$
- Unknown parameters $(\mu_k, \Sigma_k)$ and $\pi_k$, with $\pi_k$ satisfying the condition $\sum_{k=1}^{3} \pi_k = 1$.

We will assign IF values, $x_{ij}$ in a given genomic distance $D_j$ using hierarchical model-based clustering. Briefly, given an initial beliefs about $z_{ij}$, which represents the indicator of the cluster for a IF value $x_{ij}$, and parameters $(\pi, \mu_{1:3}, \Sigma_{1:3})$, the probability of $x_{ij}$ under a cluster $k$ is assumed to be:

$$p(z_{ij} = k | x_{ij}) = \frac{p(z_{ij} = k) p(x_{ij} | z_{ij} = k)}{p(x_{ij})} = \frac{\pi_k \phi(x_{ij}; \mu_k, \Sigma_k)}{\sum_{l=1}^{3} \pi_l \, \phi(x_{ij}; \mu_l, \Sigma_l)}$$

where:

- $p(z_{ij} = k)$ is the prior probability that $x_{ij}$ belongs to cluster $k$, with prior weight $\pi_k$,
- $p(x_{ij} | z_{ij} = k)$ is the probability of observing $x_{ij}$ under cluster $k$.

Prior to observing $x_{ij}$, we assume a prior probability $\pi_k$ that $x_{ij}$ belongs to cluster $k$. Upon observing $x_{ij}$, we update this belief based on the likelihood of $x_{ij}$ under each cluster. This entire process of estimating the unknown parameters and $p(z_{ij} = k | x_{ij})$ is optimized by the Expectation-Maximization (EM) algorithm.

The assignment of an IF value to a specific cluster is determined by selecting the cluster that maximizes the conditional probability, given the estimated parameters:

$$\text{argmax}_k \, p(z_{ij} = k | x_{ij})$$

**Supplemental Note S4. Consistent test of different number of single cell in esach group**

To evaluate the consistency of the scHiCcompare workflow under varying numbers of scHi-C data in each group (imbalanced scenarios), we designed a controlled experiment. Using 600 scHi-C data samples from ODC cell types with a known ground truth, we created two condition groups with 300 samples each. From these, we generated subsets representing imbalanced cases (300 vs. 200, 300 vs. 150, and 300 vs. 100 samples per group). We then applied scHiCcompare with progressive pooling and Random Forest imputation to examine performance across these setups (each case is run with 20 iterations).

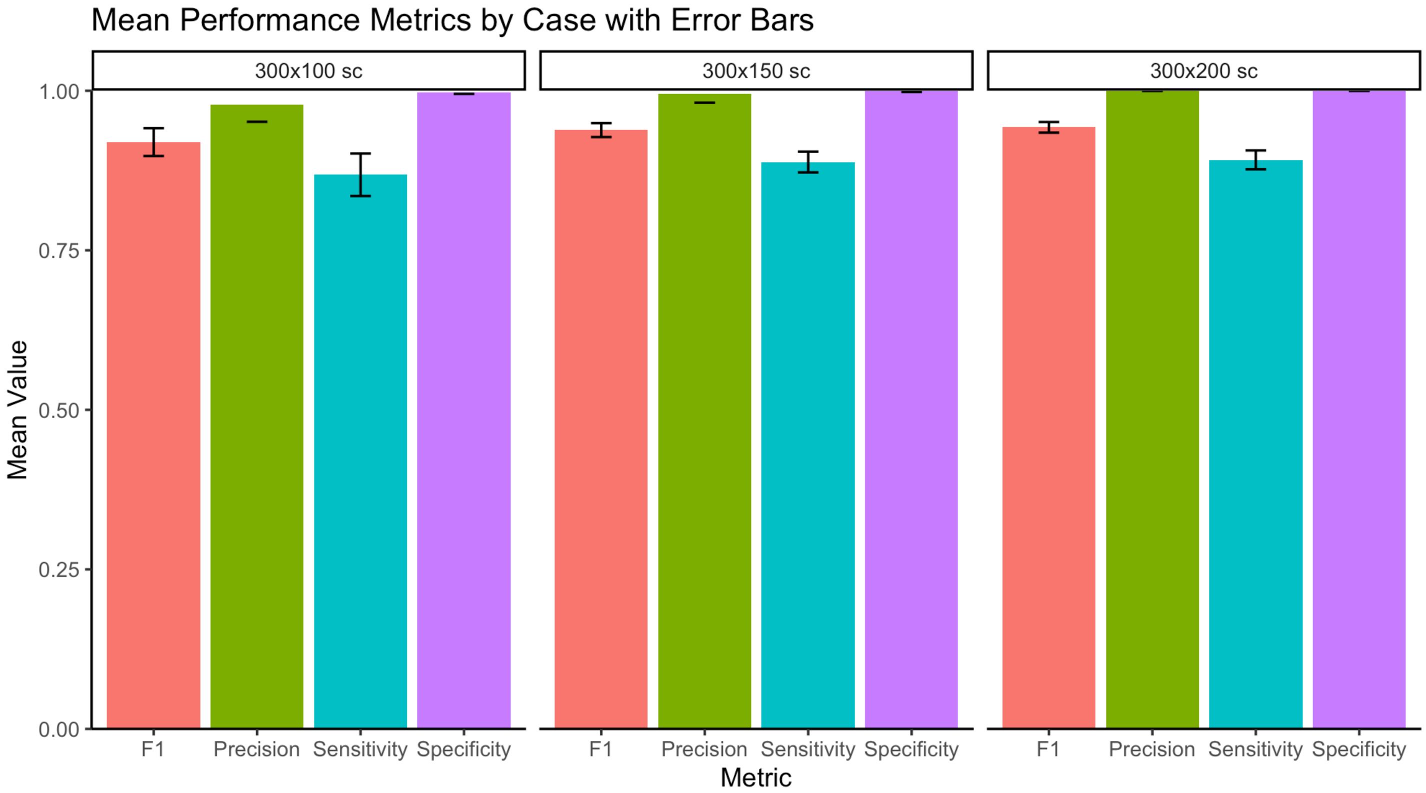


**Figure S1. Comparison of the scHiCcompare performance over different number of single-cell in each group.** These barplots displays the values of mean for four performance metrics—F1, Precision, Sensitivity, and Specificity—across three different cases labeled as 300x100 sc, 300x150 sc, and 300x200 sc. The error bars indicate variability for each metric in each case.

The result showed that the scHiCcompare performs consistently well across different cases, achieving high values for F1, Precision, Sensitivity, and Specificity. The small error bars indicate small variability or changes, suggesting that the workflow's performance is stable and robust across different sample sizes in each group.

**Supplemental Figure S2 and Supplemental Figure S3 Imputed values distribution compares with observation data**

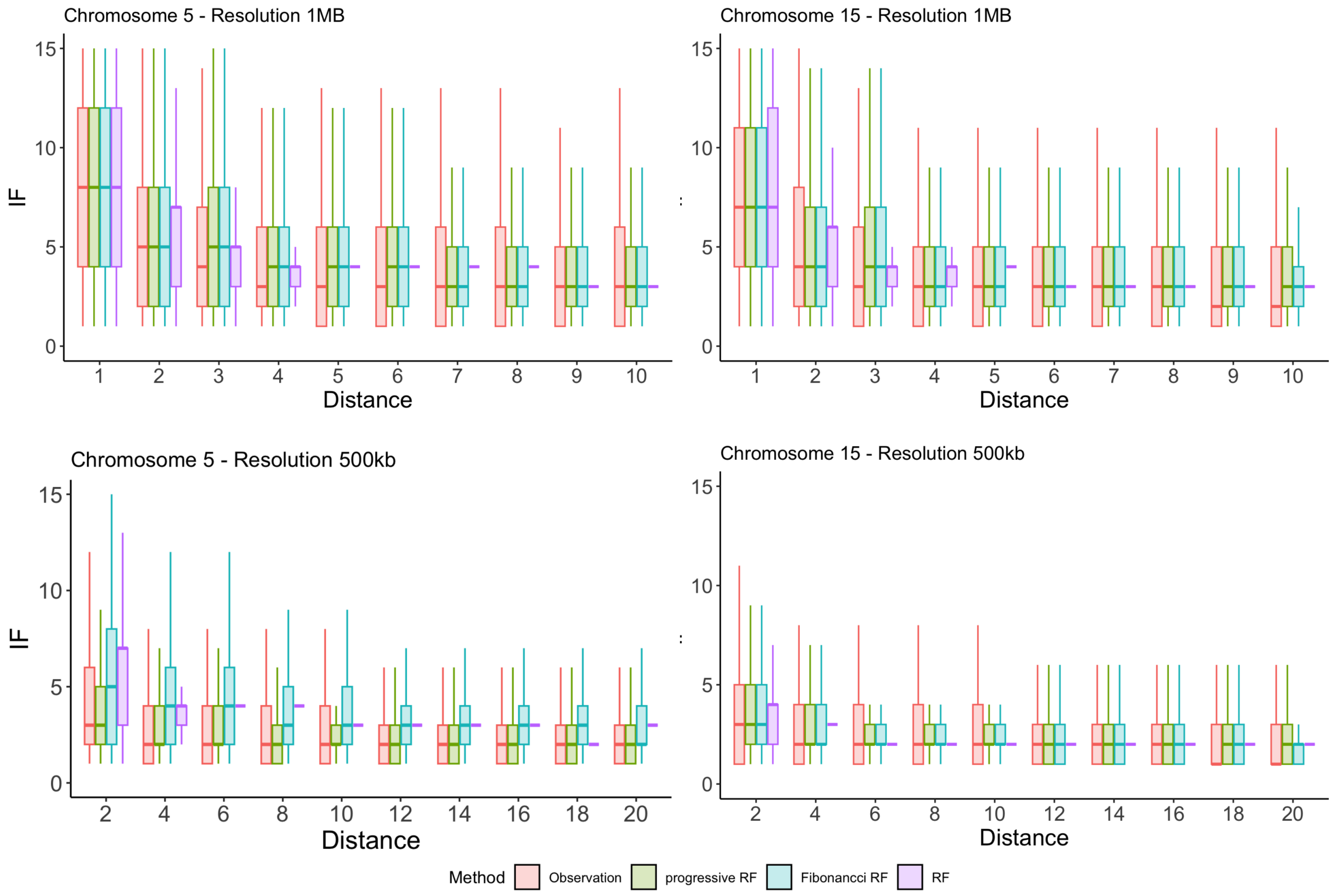


This boxplot illustrates the range and interquartile range (IQR) of observed and imputed Imputation Fidelity (IF) values for a set of 100 oocyte-derived cell (ODC) single-cell matrices across various genomic distances (ranging from 1MB to 10MB) for each imputation method. The imputation strategies include standard Random Forest (RF), progressive RF pooling, and Fibonacci RF pooling. The x-axis represents genomic distance, scaled according to the resolution used (1MB or 500kb).

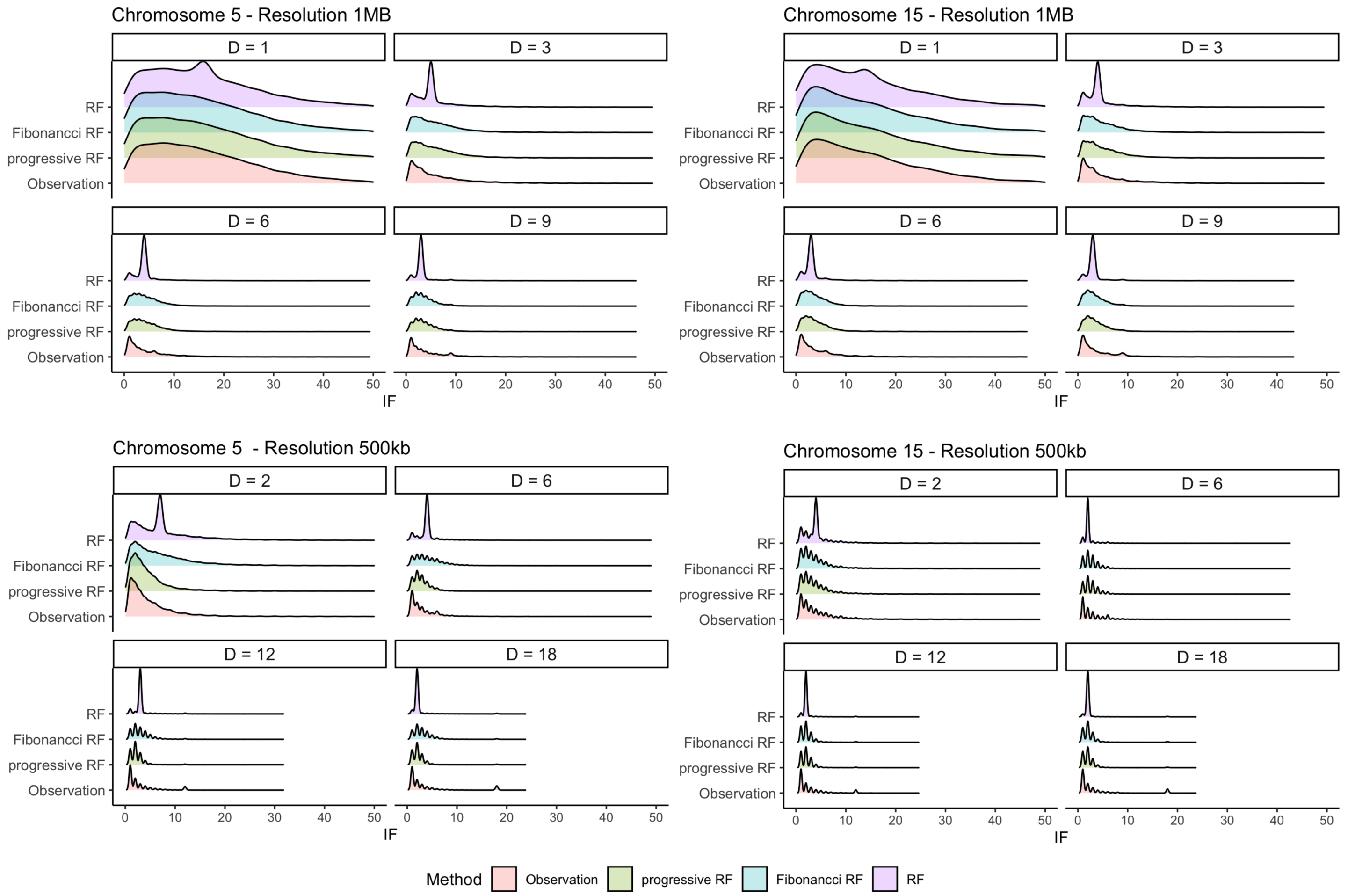


This figure shows the distribution of observed and imputed IF values for a group of 100 oocyte-derived cells (ODCs) single-cell matrices. The imputation was performed using different Random Forest strategies: standard Random Forest (RF), Random Forest with progressive pooling (progressive RF), and Random Forest with Fibonacci-style pooling (Fibonacci RF). The distributions are displayed for selected genomic distances (1MB, 3MB, 6MB, and 9MB).

**Supplemental Figure S4. Other measurements in comparision of normalization methods**

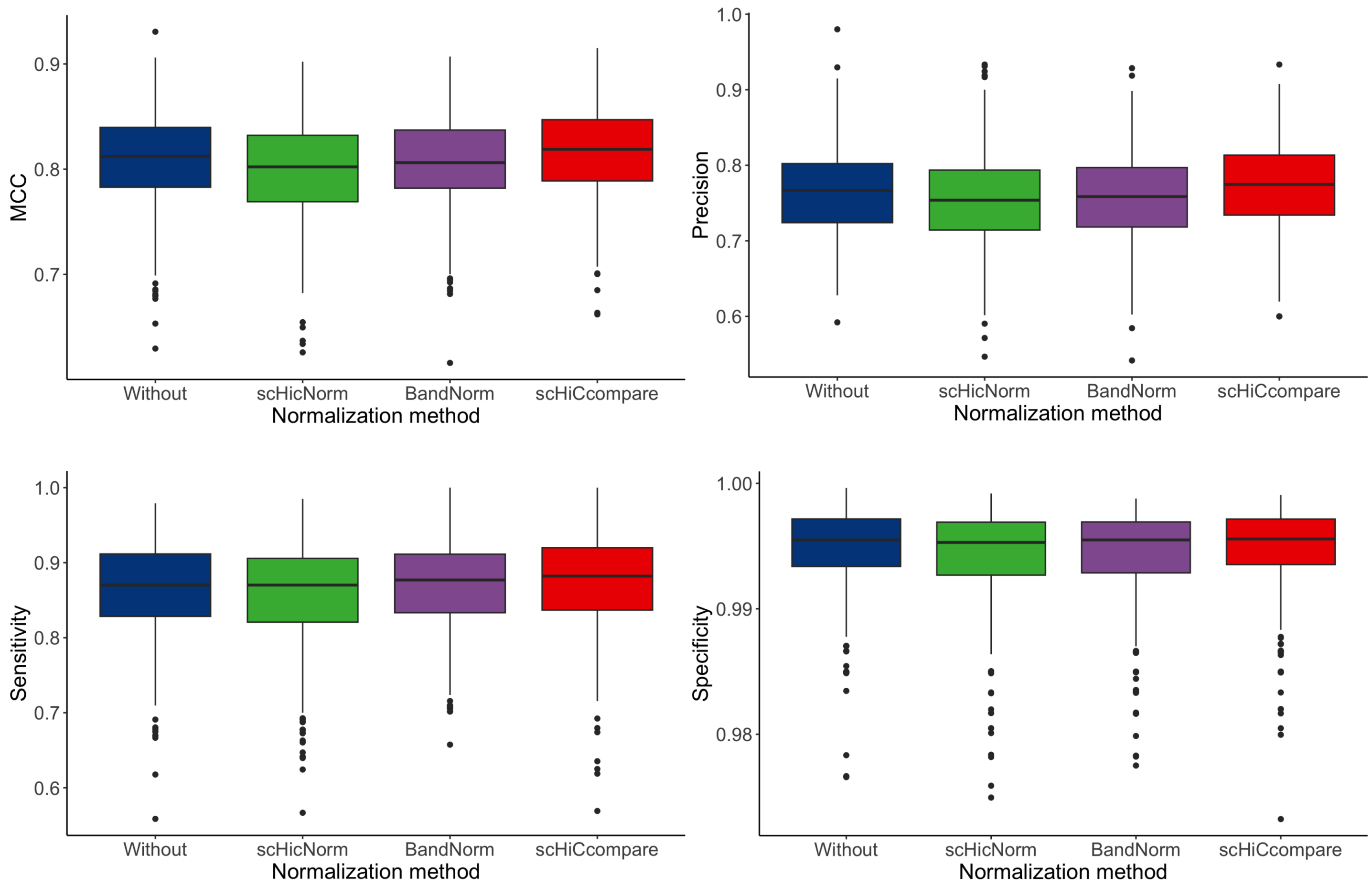


This figure shows the performance comparison of four normalization methods—Without normalization (Before), scHiCcompare, scHiCNorm, and BandNorm—applied to controlled changes data within the scHiCcompare workflow. The methods are evaluated across four metrics: Matthews Correlation Coefficient (MCC), Precision, Sensitivity, and Specificity. Each boxplot shows the interquartile range (IQR) of metric values for each method across 22 chromosomes (each has 20 iterations)

**Supplemental Figure S5. Other measurements in comparision of differential detection methods**

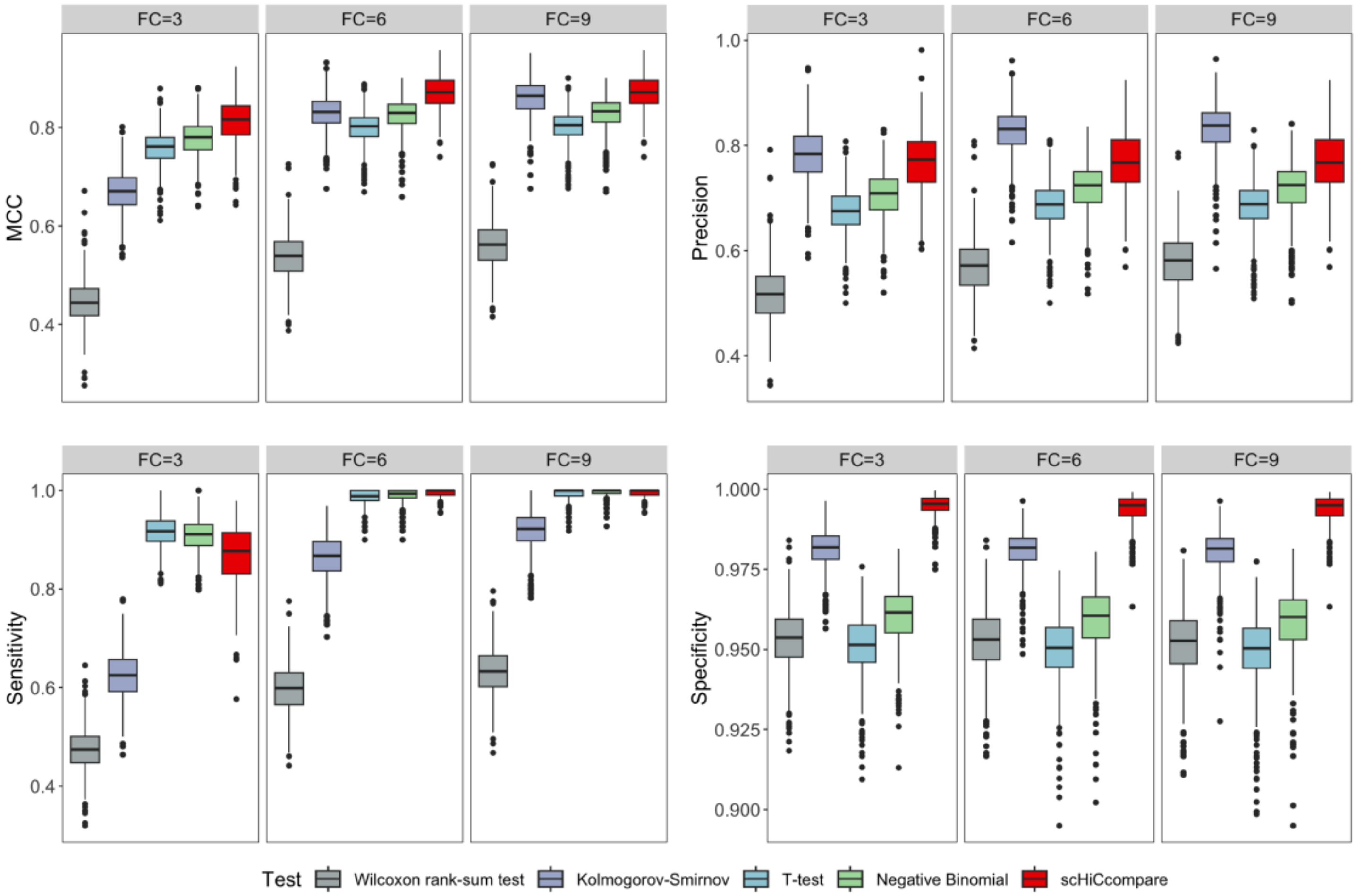


This figure shows the performance comparison of four normalization methods—Without normalization (Before), scHiCcompare, scHiCNorm, and BandNorm—applied to controlled changes data within the scHiCcompare workflow. The methods are evaluated across four metrics: Matthews Correlation Coefficient (MCC), Precision, Sensitivity, and Specificity. Each boxplot shows the interquartile range (IQR) of metric values for each method across 22 chromosomes (each has 20 iterations)

**Supplemental Figure S6. Other measurements in comparision of published workflow**

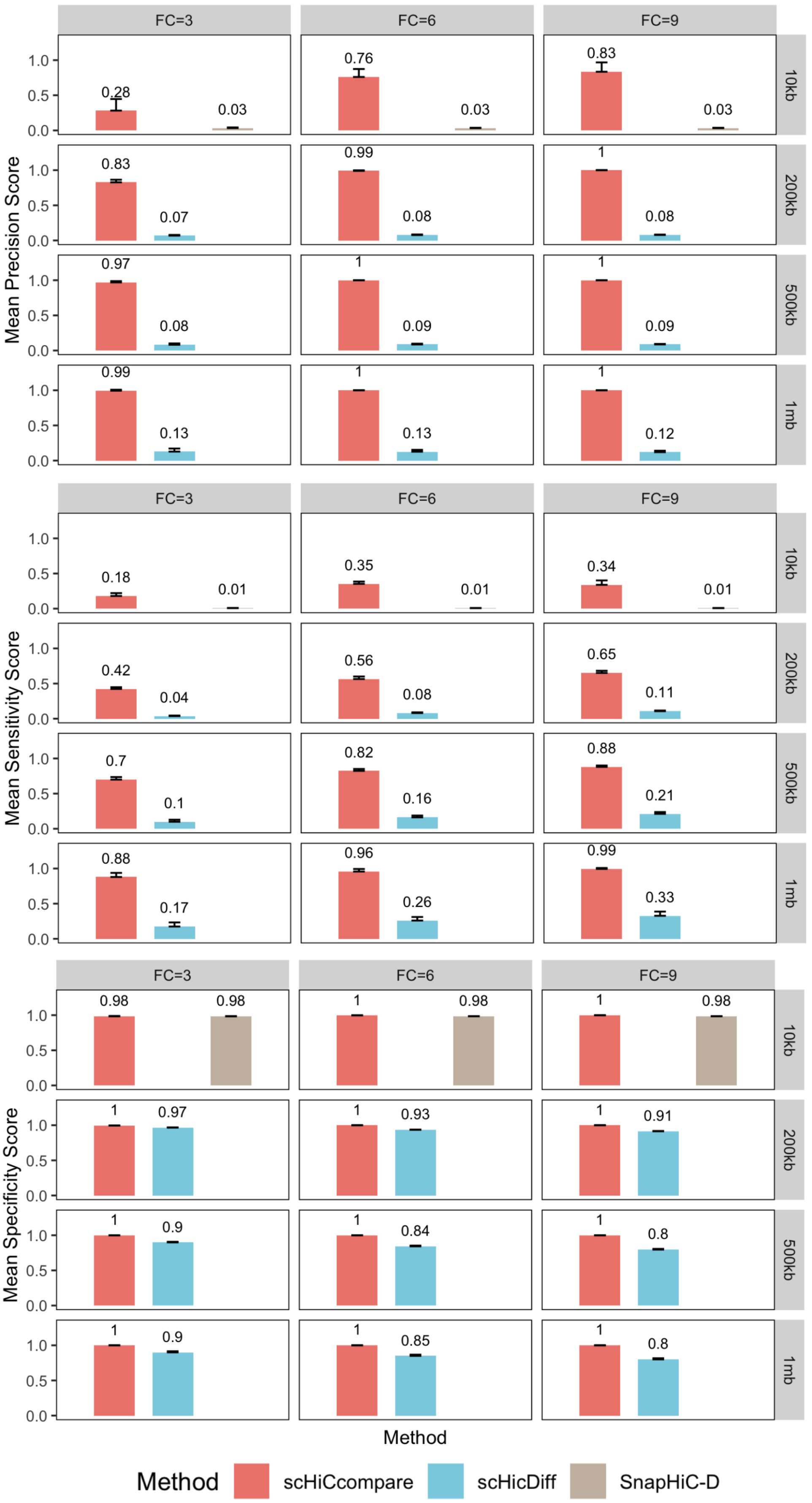

This figure compares the performance of three differential chromatin interaction detection workflow —scHiCcompare, scHiCDiff, and SnapHiC-D —across different DCI's fold change levels (FC = 3, 6, and 9) and data resolutions (10kb, 200kb, 500kb, and 1MB). The methods are evaluated on three key metrics: Precision, Sensitivity, and Specificity.

**Supplemental Figure S7. Histone and ChIP-seq peak comparison between astrocytes (Astro) and endothelial cells (Endo)**

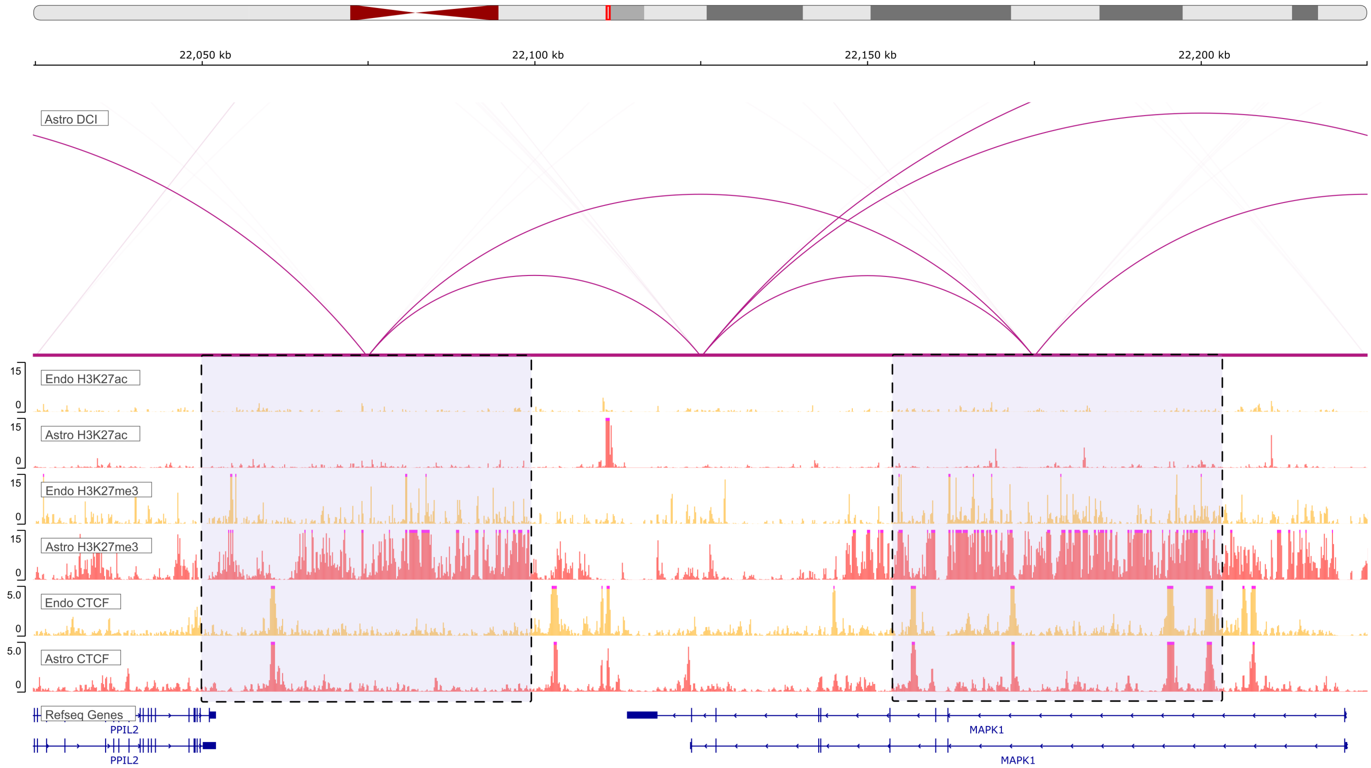


The arc track displays a sample region of differential chromatin interaction (DCI) in chromosome 22 between astrocytes and endothelial cells, where the adjusted interaction frequency is higher in astrocytes. The signal tracks display H3K27ac, H3K27me3, and CTCF signals for endothelial cells and astrocytes, in that order from top to bottom. In two purple regions of significant detected differences, the H3K27ac track in astrocytes has greater signal peak compared to that of endocytes.

**Supplemental Table S1. scHiCcompare application reveals biology of excitatory cell**

| Cell1 vs Cell2 | bulk IF1 | bulk IF2 | M | normalized bulk IF1 | normalized bulk IF2 | adj.M | Z | Difference |
|---|---|---|---|---|---|---|---|---|
| Vip vs L23 | 2,522 | 825 | -1.61 | 1,470.89 | 1,414.55 | -0.06 | -0.47 | 0 |
| Vip vs L4 | 3,136 | 3,840 | 0.29 | 3,178.53 | 3,788.62 | 0.25 | 2.19 | 1 |
| Vip vs L5 | 4,584 | 4,235 | -0.11 | 4,131.49 | 4,698.84 | 0.19 | 1.83 | 1 |
| Vip vs L6 | 898 | 1,193 | 0.41 | 954.46 | 1,122.43 | 0.23 | 1.99 | 1 |
| Ndnf vs L23 | 3,288 | 1,224 | -1.43 | 2,793.45 | 1,440.70 | -0.96 | -3.40 | 1 |
| Ndnf vs L4 | 2,776 | 3,840 | 0.47 | 2,993.82 | 3,560.61 | 0.25 | 2.16 | 1 |
| Ndnf vs L5 | 3,288 | 2,950 | -0.16 | 2,951.02 | 3,286.86 | 0.16 | 1.49 | 0 |
| Ndnf vs L6 | 1,428 | 2,287 | 0.68 | 1,670.66 | 1,954.81 | 0.23 | 1.95 | 1 |

| Cell1 vs Cell2 | bulk IF1 | bulk IF2 | M | normalized bulk IF1 | normalized bulk IF2 | adj.M | Z | Difference |
|---|---|---|---|---|---|---|---|---|

M = log fold change of (IF2/IF1); adj.M = log fold change of normalized (IF2/IF1)

**Supplemental Table S2. Overlapped regions of astrocytes (Astro) and endothelial cells (Endo) with reported differential chromatin domain boundaries**

| chr | start | end |
|---|---|---|
| chr22 | 33,375,000 | 33,400,000 |
| chr22 | 45,475,000 | 45,500,000 |
| chr22 | 45,875,000 | 45,900,000 |
| chr22 | 26,325,000 | 26,350,000 |
| chr22 | 45,450,000 | 45,475,000 |
| chr22 | 48,850,000 | 48,875,000 |
| chr22 | 50,700,000 | 50,725,000 |
| chr22 | 49,050,000 | 49,075,000 |
| chr22 | 30,700,000 | 30,725,000 |
| chr22 | 17,450,000 | 17,475,000 |
| chr22 | 45,400,000 | 45,425,000 |
| chr22 | 20,050,000 | 20,075,000 |
| chr22 | 28,025,000 | 28,050,000 |
| chr22 | 45,225,000 | 45,250,000 |
| chr22 | 19,275,000 | 19,300,000 |
| chr22 | 51,100,000 | 51,125,000 |
| chr22 | 37,200,000 | 37,225,000 |
| chr22 | 19,525,000 | 19,550,000 |
| chr22 | 26,675,000 | 26,700,000 |
| chr22 | 29,525,000 | 29,550,000 |
| chr22 | 35,050,000 | 35,075,000 |
| chr22 | 39,275,000 | 39,300,000 |
| chr22 | 45,525,000 | 45,550,000 |
| chr22 | 45,550,000 | 45,575,000 |
| chr22 | 49,600,000 | 49,625,000 |

| chr | start | end |
|---|---|---|
| chr22 | 35,425,000 | 35,450,000 |
| chr22 | 25,825,000 | 25,850,000 |
| chr22 | 36,125,000 | 36,150,000 |
| chr22 | 49,450,000 | 49,475,000 |
| chr22 | 38,475,000 | 38,500,000 |
| chr22 | 20,700,000 | 20,725,000 |
| chr22 | 23,350,000 | 23,375,000 |
| chr22 | 19,500,000 | 19,525,000 |
| chr22 | 48,650,000 | 48,675,000 |

**Supplemental Table S3. Comparison of median signal (Wilcoxon p-values) for Endo and Astro histone markers and CTCF**

| Signal | Endo Histone Marker | | | Astro Histone Marker | | |
|---|---|---|---|---|---|---|
| Signal | Median of Endo | Median of Astro | p.value | Median of Endo | Median of Astro | p.value |
| H3K27ac | 3,619.65 | 3,132.74 | 0.29 | 2,959.69 | 2,995.63 | 0.38 |
| H3K27me3 | 7,254.34 | 7,453.79 | 0.47 | 4,264.23 | 6,320.15 | 0.00 |
| CTCF | 1,918.81 | 1,902.49 | 0.48 | 870.09 | 1,042.29 | 0.00 |

**Supplemental Table S4. Summary fold change of DCI between oocytes (non-surrounded nucleolus, NSN) and mature oocytes (surrounded nucleolus, SN)**

| D | Up.FC_Median | Up.FC_Mean | Up.FC_Sd | Down.FC_Median | Down.FC_Mean | Down.FC_Sd |
|---|---|---|---|---|---|---|
| 1 | 1.32 | 1.39 | 0.24 | 1.31 | 1.37 | 0.22 |
| 2 | 1.31 | 1.37 | 0.23 | 1.30 | 1.36 | 0.22 |
| 3 | 1.27 | 1.33 | 0.20 | 1.26 | 1.32 | 0.19 |
| 4 | 1.26 | 1.32 | 0.19 | 1.26 | 1.31 | 0.18 |
| 5 | 1.24 | 1.29 | 0.16 | 1.24 | 1.28 | 0.16 |
| 6 | 1.23 | 1.27 | 0.15 | 1.22 | 1.27 | 0.14 |
| 7 | 1.22 | 1.27 | 0.14 | 1.22 | 1.27 | 0.15 |
| 8 | 1.22 | 1.26 | 0.13 | 1.21 | 1.25 | 0.13 |
| 9 | 1.21 | 1.25 | 0.12 | 1.21 | 1.24 | 0.12 |
| 10 | 1.21 | 1.24 | 0.12 | 1.20 | 1.24 | 0.11 |

| D | Up.FC_Median | Up.FC_Mean | Up.FC_Sd | Down.FC_Median | Down.FC_Mean | Down.FC_Sd |
|---|---|---|---|---|---|---|
| 11 | 1.20 | 1.24 | 0.12 | 1.20 | 1.23 | 0.11 |
| 12 | 1.20 | 1.23 | 0.11 | 1.20 | 1.22 | 0.10 |
| 13 | 1.20 | 1.23 | 0.10 | 1.19 | 1.22 | 0.11 |
| 14 | 1.20 | 1.23 | 0.10 | 1.20 | 1.22 | 0.10 |
| 15 | 1.20 | 1.23 | 0.09 | 1.25 | 1.27 | 0.12 |
| 16 | 1.23 | 1.25 | 0.09 | 1.21 | 1.25 | 0.13 |
| 17 | 1.17 | 1.19 | 0.08 | 1.22 | 1.27 | 0.16 |
| 18 | 1.17 | 1.20 | 0.07 | 1.21 | 1.26 | 0.16 |
| 19 | 1.17 | 1.19 | 0.08 | 1.21 | 1.27 | 0.16 |
| 20 | 1.17 | 1.19 | 0.07 | 1.21 | 1.26 | 0.16 |
| 21 | 1.24 | 1.26 | 0.09 | 1.24 | 1.27 | 0.11 |
| 22 | 1.21 | 1.24 | 0.10 | 1.24 | 1.26 | 0.11 |
| 23 | 1.19 | 1.22 | 0.11 | 1.21 | 1.25 | 0.14 |
| 24 | 1.18 | 1.22 | 0.11 | 1.22 | 1.27 | 0.16 |
| 25 | 1.19 | 1.23 | 0.10 | 1.21 | 1.26 | 0.15 |
| 26 | 1.19 | 1.22 | 0.10 | 1.20 | 1.25 | 0.14 |
| 27 | 1.18 | 1.21 | 0.10 | 1.21 | 1.26 | 0.14 |
| 28 | 1.22 | 1.24 | 0.09 | 1.20 | 1.25 | 0.13 |
| 29 | 1.19 | 1.23 | 0.12 | 1.22 | 1.24 | 0.11 |
| 30 | 1.19 | 1.22 | 0.11 | 1.20 | 1.24 | 0.13 |
| 31 | 1.19 | 1.21 | 0.10 | 1.22 | 1.25 | 0.15 |
| 32 | 1.19 | 1.21 | 0.11 | 1.22 | 1.25 | 0.14 |
| 33 | 1.19 | 1.22 | 0.11 | 1.22 | 1.25 | 0.15 |
| 34 | 1.19 | 1.22 | 0.10 | 1.21 | 1.25 | 0.15 |
| 35 | 1.19 | 1.23 | 0.12 | 1.20 | 1.25 | 0.15 |
| 36 | 1.19 | 1.22 | 0.12 | 1.21 | 1.25 | 0.15 |
| 37 | 1.18 | 1.22 | 0.12 | 1.22 | 1.25 | 0.13 |
| 38 | 1.18 | 1.22 | 0.12 | 1.21 | 1.25 | 0.15 |
| 39 | 1.19 | 1.22 | 0.11 | 1.22 | 1.24 | 0.13 |
| 40 | 1.17 | 1.21 | 0.11 | 1.21 | 1.23 | 0.12 |
| 41 | 1.17 | 1.21 | 0.10 | 1.19 | 1.23 | 0.13 |

| D | Up.FC_Median | Up.FC_Mean | Up.FC_Sd | Down.FC_Median | Down.FC_Mean | Down.FC_Sd |
|---|---|---|---|---|---|---|
| 42 | 1.17 | 1.20 | 0.09 | 1.21 | 1.24 | 0.13 |
| 43 | 1.17 | 1.21 | 0.09 | 1.20 | 1.24 | 0.12 |
| 44 | 1.18 | 1.21 | 0.10 | 1.19 | 1.24 | 0.13 |
| 45 | 1.17 | 1.20 | 0.09 | 1.21 | 1.24 | 0.12 |
| 46 | 1.18 | 1.21 | 0.09 | 1.20 | 1.23 | 0.11 |
| 47 | 1.18 | 1.21 | 0.09 | 1.19 | 1.23 | 0.14 |
| 48 | 1.18 | 1.21 | 0.10 | 1.21 | 1.23 | 0.13 |
| 49 | 1.17 | 1.20 | 0.09 | 1.21 | 1.24 | 0.13 |
| 50 | 1.18 | 1.20 | 0.08 | 1.20 | 1.23 | 0.14 |

**Supplemental Table S5. Number of detected differential chromatin interaction (DCI) between oocytes (non-surrounded nucleolus, NSN) and mature oocytes (surrounded nucleolus, SN)**

| Chr | n_up.regulation | n_down.regulation |
|---|---|---|
| 1 | 7,673 | 6,666 |
| 2 | 6,901 | 5,883 |
| 3 | 5,892 | 5,639 |
| 4 | 5,252 | 4,831 |
| 5 | 5,147 | 4,531 |
| 6 | 4,933 | 4,878 |
| 7 | 5,143 | 4,655 |
| 8 | 3,796 | 3,642 |
| 9 | 3,601 | 3,311 |
| 10 | 3,906 | 3,827 |
| 11 | 3,642 | 3,299 |
| 12 | 3,705 | 3,352 |
| 13 | 3,762 | 3,341 |
| 14 | 3,500 | 3,286 |
| 15 | 2,793 | 2,759 |
| 16 | 2,582 | 2,621 |
| 17 | 2,063 | 1,999 |

| Chr | n_up.regulation | n_down.regulation |
|---|---|---|
| 18 | 2,230 | 2,123 |
| 19 | 845 | 879 |

**Supplemental Table S6. Running time of scHiCcompare, SnapHiC-D, and scHiCDiff workflow in different resolutions (second unit)**

| Resolution | Sc_N | ScHiCcompare_RF | ScHiCcompare_RF + progress | ScHiCcompare_RF + Fibbonancci | ScHiCDiff | SnapHiC-D |
|---|---|---|---|---|---|---|
| 10kb | 100 | 27,855.380 | 32,107.600 | 24,361.190 | | 37,440 |
| 200kb | 100 | 21.787 | 93.600 | 91.144 | 2,250.70500 | |
| 500kb | 100 | 6.135 | 26.620 | 28.264 | 26.15860 | |
| 1MB | 100 | 2.870 | 8.087 | 9.478 | 15.70075 | |

Note: SnapHiC-D only works on 10kb resolution data. Due to excessive memory and long run time, scHiCDiff was not tested for 10kb resolution.

### Reference

[1] L.L. Doove, S. Van Buuren, E. Dusseldorp, Recursive partitioning for missing data imputation in the presence of interaction effects, Computational Statistics & Data Analysis 72 (2014) 92–104.
